# Shield-4i: A Whole-mount Multiplexed Imaging Platform for Studying Multiscale Information Flow in 3D Multicellular Systems

**DOI:** 10.64898/2026.07.01.735871

**Authors:** Ruth Hornbachner, Shayan Shamipour, Feyza Nur Arslan, Rui Fan, Max Hess, Florian Curvaia, Joel Lüthi, Andrew C. Oates, Ivan Bedzhov, Darren Gilmour, Virginie Uhlmann, Lucas Pelkmans

## Abstract

Self-organization in multicellular systems emerges from reciprocal interactions across spatiotemporal scales. Understanding how subcellular organization, tissue remodeling and developmental outcome are coordinated, thus requires simultaneous profiling of biological processes spanning orders of magnitudes in space and time. Yet, a unified experimental and computational framework for capturing these multiscale properties across *in vivo* and stem cell-derived systems has been lacking. Here, we introduce **Shield-4i**, a high-throughput, versatile, and accessible method for automated *in toto* iterative immunofluorescence imaging of whole-mount structures at subcellular resolution. Through polyepoxide-mediated inter- and intramolecular crosslinking, Shield-4i preserves sample integrity during repeated SDS-based elution cycles. We benchmark this method in gastrulating zebrafish and post-implantation mouse embryos and demonstrate its applicability to stem cell-derived 3D gastruloids, achieving up to 30-plex measurements of proteins and their post-translational modifications across hundreds of samples. To enable scalable analysis, we developed a dedicated 3D workflow supporting OME-Zarr-based and FAIR-compliant data storage, standardized processing, and multiscale feature extraction. Applying this framework to investigate gastruloid self-organization, we quantify how cellular physicochemical state and signaling properties encode cell position along embryonic axes and connect molecular patterning and fate decisions to morphological symmetry breaking at the multicellular scale. Together, Shield-4i provides a high-content *in toto* spatial proteomics platform for dissecting multiscale information flow and self-organization in multicellular systems.

## Introduction

Starting from a single-cell zygote or stem cells in culture, multicellular self-organization emerges through scale-crossing interactions between molecular, cellular, and tissue-level processes, collectively shaping developmental outcomes^1,2^. Local processes such as cytoskeletal remodeling or cell-extracellular matrix interactions can reshape tissue architecture, while the resulting geometry and mechanics feed back on signaling, adhesion and fate decisions^3–8^. These reciprocal interactions distribute information across scales, with subcellular organization, cellular decisions and tissue morphology jointly controlling the developmental outcome. Understanding how such scale-crossing information flows are established, maintained, and transformed into reproducible trajectories, thus requires experimental and computational frameworks that connect the molecular phenotype of individual cells to their neighbourhood composition and global tissue architecture.

Proteins, their activity states, and the higher-order assemblies they form are particularly well suited for such scale-crossing measurements, as they integrate diverse signaling inputs into the physicochemical state of the cell - the integrated biochemical and mechanical state that governs cellular behaviour - and translate these into contextual cellular responses^9^. Advances in spatial proteomics have enabled highly multiplexed mapping of protein and protein state distributions across diverse tissue types, revealing cell-type composition, signaling dynamics, and the architecture of complex tissue microenvironments^10,11^. However, most spatial multiplexing approaches remain restricted to two-dimensional cultures or thin tissue sections^12–14^. While we recently achieved up to ten protein state measurements in whole-mount zebrafish embryos^15^, highly multiplexed profiling of three-dimensional specimens remains challenging, due to loss of antigenicity and sample deterioration during multiple staining and elution cycles. Crosslinking strategies that stabilize RNAs, proteins, and tissue structure during harsh delipidation steps provide a route to address this limitation, highlighted by their successful use in tissue clearing and multiplex protein measurements in tissue blocks^16–19^. Yet, the integration of these methods into a scalable platform for high-dimensional whole-mount imaging across developmental systems and the resulting multiscale analysis remains unresolved.

To address these challenges, we integrate SHIELD-based stabilization through intramolecular epoxide linkages^20^ with iterative indirect immunofluorescence imaging (4i)^12^ that employs high-concentration, high-temperature SDS for antibody elution and tissue clearing. This strategy, termed Shield-4i, achieves uniform staining and elution while preserving antigenicity and sample structure up to 24 imaging cycles across 90 days. Combined with robust immobilization, optical clearing, gentle liquid handling, and high-throughput spinning-disk confocal microscopy, this approach enables automated highly multiplexed volumetric imaging of hundreds of samples. We applied Shield-4i to gastrulating zebrafish embryos, post-implantation mouse embryos, and stem cell-derived murine gastruloids up to 350 µm in thickness, acquiring up to 30-plex *in toto* protein abundance and state measurements and demonstrating its versatility across model systems with distinct architectures and cell densities.

Importantly, to use the resulting terabyte-scale multidimensional datasets in quantitative and systematic studies that integrate measurements from large numbers of multicellular structures, we developed a generalizable computational task collection for 3D multiplexed imaging datasets, encompassing file conversion, illumination correction, image registration, and multiscale object segmentation and feature extraction, that can be applied to diverse structures and tissue types. This task collection runs in Fractal, an open-source no-code workflow orchestration platform for large-scale microscopy analysis on high-performance computing clusters that we recently developed^21^, which relies on the next-generation file format OME-Zarr^22^ for Findable, Accessible, Interoperable, and Reusable (FAIR) high-content microscopy data^23^. Collectively, Shield-4i provides a comprehensive framework for highly multiplexed molecular profiling of 3D structures at subcellular resolution across hundreds of samples from both *in vivo* and stem cell-derived embryo systems, while supporting integrated, scalable, and tractable analysis workflows.

## Results

### Establishing Shield-4i to extend multivariate protein measurements in 3D tissues

To achieve highly multiplexed protein measurements in 3D structures using Shield-4i, PFA-fixed samples were immobilized at the bottom of Poly-D-Lysine/Fibronectin-coated 96-well plates using laser-induced crosslinking, as previously described^15^ (Methods). Combined with gentle liquid exchange via automated pipetting, this approach reliably stabilized 3D structures during iterative staining and imaging, achieving high retention efficiency across consecutive staining cycles (Supp Fig. 1a-b). Samples were subsequently protected using epoxy and Shield buffers using a protocol adapted from one previously applied on brain tissue blocks^20^. Briefly, co-incubation of epoxy with Shield buffers at 4°C and pH 7.4 allows its homogeneous diffusion throughout the tissue, and a shift to 37°C and pH 10.0 triggers uniform inter- and intramolecular crosslinking. This process preserves nucleic acids, protein antigenicity, and tissue architecture during subsequent harsh elution steps^20^. To clear the tissue and/or minimize residual signal from the previous staining cycle, we applied an elution step using 8% SDS at 70 °C for 2 hours at each cycle. This was followed by extensive washing in 1% PBS-Triton X-100 for several hours to overnight, with incubation times optimized for each tissue type at 37°C to ensure the complete removal of residual SDS, as remaining SDS strongly impairs antigen-antibody binding. Following clearing and permeabilization, samples were processed for indirect immunofluorescence imaging (Supp Fig. 1a, Methods).

To assess the multiplexing performance of Shield-4i in whole-mount tissues, we used gastrulating zebrafish embryos as a benchmark and subjected them to repeated cycles of immunofluorescence staining, elution and re-staining (Fig. 1a-b). Using nuclear (PCNA) and membrane (E-Cadherin) markers, we evaluated i) preservation of tissue morphology, ii) efficiency of antibody signal removal after elution, and iii) maintenance of antigenicity upon re-staining. We found that samples imaged across different imaging cycles deformed only mildly (Supp Fig. 1c) and could be robustly registered to a reference cycle using the registration pipelines discussed below^24,25^, achieving a median single-cell Pearson’s correlation coefficient of 0.98 for nuclear DAPI intensity (Fig. 1d and Supp Fig. 1d). This high correspondence indicates that cycle-to-cycle deformations are sufficiently small to be effectively corrected using image registration algorithms. Moreover, the antibody signals reduced to background levels following consecutive elution steps, and were largely recovered upon restaining with the same antibodies, displaying high single-cell intensity correlations with the initial staining (Fig. 1c-e). These results demonstrate that Shield-based crosslinking effectively preserves tissue integrity and protein antigenicity, while high-temperature SDS clearing enables efficient signal removal. This combination supports consecutive rounds of immunofluorescence imaging in whole-mount structures without a gradual accumulation of signal from non-eluted antibodies, allowing for highly reproducible and quantitative multiplexed protein measurements therein.

**Fig. 1.**
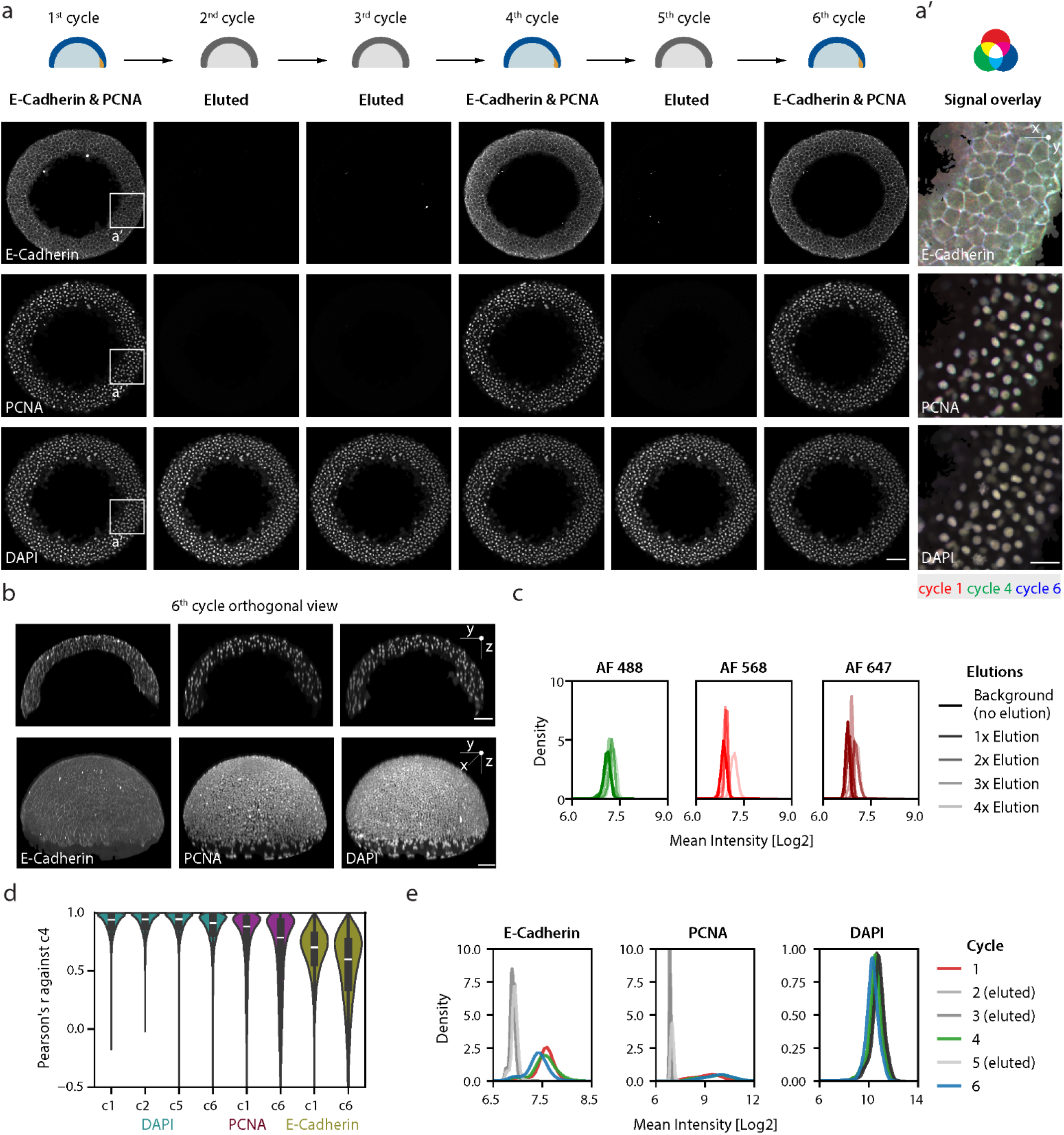
Shield-4i ensures efficient signal removal, while preserving antigenicity across multiple cycles of immunofluorescence imaging. a. Representative 2D cross-sections of a gastrulating zebrafish embryo at 5.7 hours post fertilization stained for E-Cadherin, PCNA and DAPI across consecutive Shield-4i elution and re-staining cycles. Cycles 2, 3 and 5 correspond to elution controls, in which primary antibodies were omitted. Cycles 4 and 6 show re-staining of the same markers acquired in the same channels as in cycle 1. The ROIs indicate the representative embryo shown in a’. Scale bar, 75 μm. a’. Zoom-ins of E-Cadherin, PCNA and DAPI stainings and re-staining patterns for the embryo shown in a. Scale bar, 20 μm. b. Orthogonal views (top) and 3D rendering (bottom) of the E-Cadherin, PCNA and DAPI stainings in the 6th imaging cycle of the embryo shown in a. Scale bars, 75 μm. c. Density plots of mean nuclear intensities for each fluorescent channel (AF: Alexa Fluor) measured in control wells lacking primary antibodies at first cycle, representing background before elution, and after 1 to 4 elution cycles, demonstrating the absence of background accumulation across repeated cycles. d. Pearson’s pixel-wise correlation coefficient per cell for membrane E-Cadherin or nuclear PCNA and DAPI intensities across staining cycles 1 to 6, when the stain was present, calculated relative to the 4th imaging cycle used as reference. e. Density plots of mean membrane E-Cadherin intensity, nuclear PCNA and DAPI intensities across staining cycles 1 to 6.

### Shield-4i enables highly multiplexed whole-mount imaging of vertebrate embryos and gastruloids

Next, we assessed the applicability of Shield-4i for multiplexed protein profiling across diverse 3D developmental and stem cell-derived systems spanning a broad range of tissue thicknesses, nuclear densities, and morphological complexities. Given the central role of gastrulation as a key developmental stage in embryonic development, marked by extensive morphogen patterning, cell fate specification and tissue remodeling^26^, we focused on gastrulating zebrafish embryos (5.7-6 hours post fertilization), post-implantation mouse embryos (embryonic E5.5 - E6.5), and murine embryonic stem cell-derived gastruloids fixed at 5 days after seeding (representing an equivalent stage to E8.5 mouse embryos^27,28^). Gastruloids were generated from either 100 or 300 starting cells, to further probe the generalizability of the method.

We assembled a broad antibody panel, capturing molecular processes underlying patterning, lineage specification and cellular physicochemical states. To monitor germ layer segregation and regional identity during gastrulation, we included lineage-associated transcription factors such as Otx2 and Nanog, Brachyury, Gata6, Sox17, Sox2, and AP2γ marking anterior epiblast, primitive streak, cardiac mesoderm, primitive endoderm, epiblast/neural and trophoblast/primordial germ cell lineages, respectively^29–35^. Because cell fate decisions are coordinated by spatially organized morphogen signals, we also included downstream readouts of major signaling pathways, including pERK, Smad 2/3, pSmad 1/5, and β-Catenin as effectors of FGF, TGF-β, and canonical Wnt signaling^36–38^. Finally, given the strong modulating effect of the physicochemical state of a cell on information processing and decision making in response to extracellular growth factors^9,39^, we also included antibodies against proteins reporting on cell architecture, mechanical state, transcriptional activity, proliferation, membrane trafficking, intracellular organization, and metabolism. These comprised cytoskeletal and mechanotransductive proteins, including E- and N-Cadherin, pMyosin, pERM, ZO1, Tubulin, Keratins, and Yap1; transcriptional and chromatin-state readouts, including Pol II S2p, Pol II S5p, H3K9me3, and H3K27ac; proliferation and apoptosis-associated proteins, including pH3, PCNA, Ki67 and cleaved caspase-3; endomembrane-associated proteins, including GM130, Cathepsin D and VPS35, marking the Golgi complex and endosomes, respectively; and the metabolic readout GLUT1. The complete list of primary and secondary antibodies used in this study is provided in Tables 1-4 (Methods). Together, this panel enables a comprehensive profiling of features of the cellular physicochemical state, signaling activities, and cell fates within the same specimen. We applied tissue-specific antibody panels to 68 zebrafish gastrulae, 75 murine gastruloids and 112 post-implantation mouse embryos, achieving 12- to 30-plex *in toto* protein state stainings (Fig. 2 and Supp. Fig. 2). Notably, we detected little-to-no measurable sample degradation or loss of antigenicity across up to 24 imaging cycles performed during 90 days. In all systems, nuclear, membrane, and cytoplasmic markers exhibited the expected subcellular localization patterns. In addition, apico-basal polarity markers (ZO1, pERM, and Pard6b) localized to the expected apical domain^40–42^ (Supp. Fig. 2b), while lineage-associated transcription factors displayed the expected spatial distributions along the anterior-posterior and proximal-distal axes of the structures^30,38,43^ (Fig. 2 and Supp. Fig. 2). We further recapitulated known signaling distributions, including elevated pERK levels in trophectoderm regions^44^ and posteriorly enriched pSmad 1/5 in mouse gastrulae^45^ (Supp. Fig. 2c). GLUT1 levels were enriched at the posterior side of gastruloids (Fig. 2b-b’’ and Supp. Fig. 2a-a’), consistent with recent findings that elevated glycolytic activity supports mesendoderm fate acquisition and maintenance at the posterior pole^39,46^. In addition, we detected a differential enrichment of the mechanosensor Yap1 in both mammalian systems between anterior and posterior poles, suggesting a potential coupling between metabolic, signaling, and mechanotransductive states during anterior-posterior axis specification (Fig. 2b-c’’ and Supp. Fig. 2c). Although future work will be required to resolve any causal links between these processes, our findings demonstrate that Shield-4i is broadly applicable across diverse 3D developmental model systems.

**Table 1:**
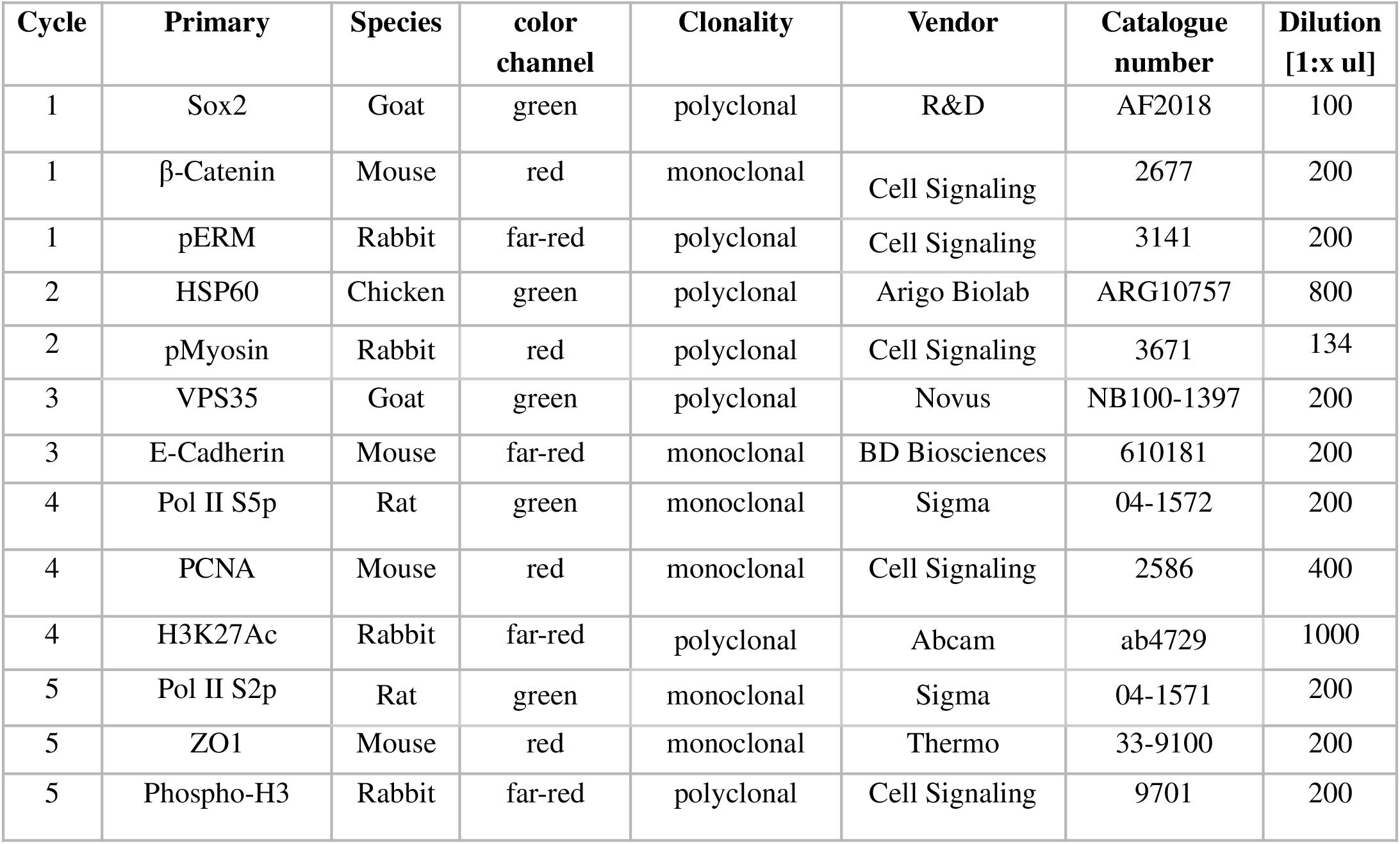
List of primary antibodies used in the multiplexing experiment on zebrafish embryos, their dilutions and staining cycles.

**Fig. 2.**
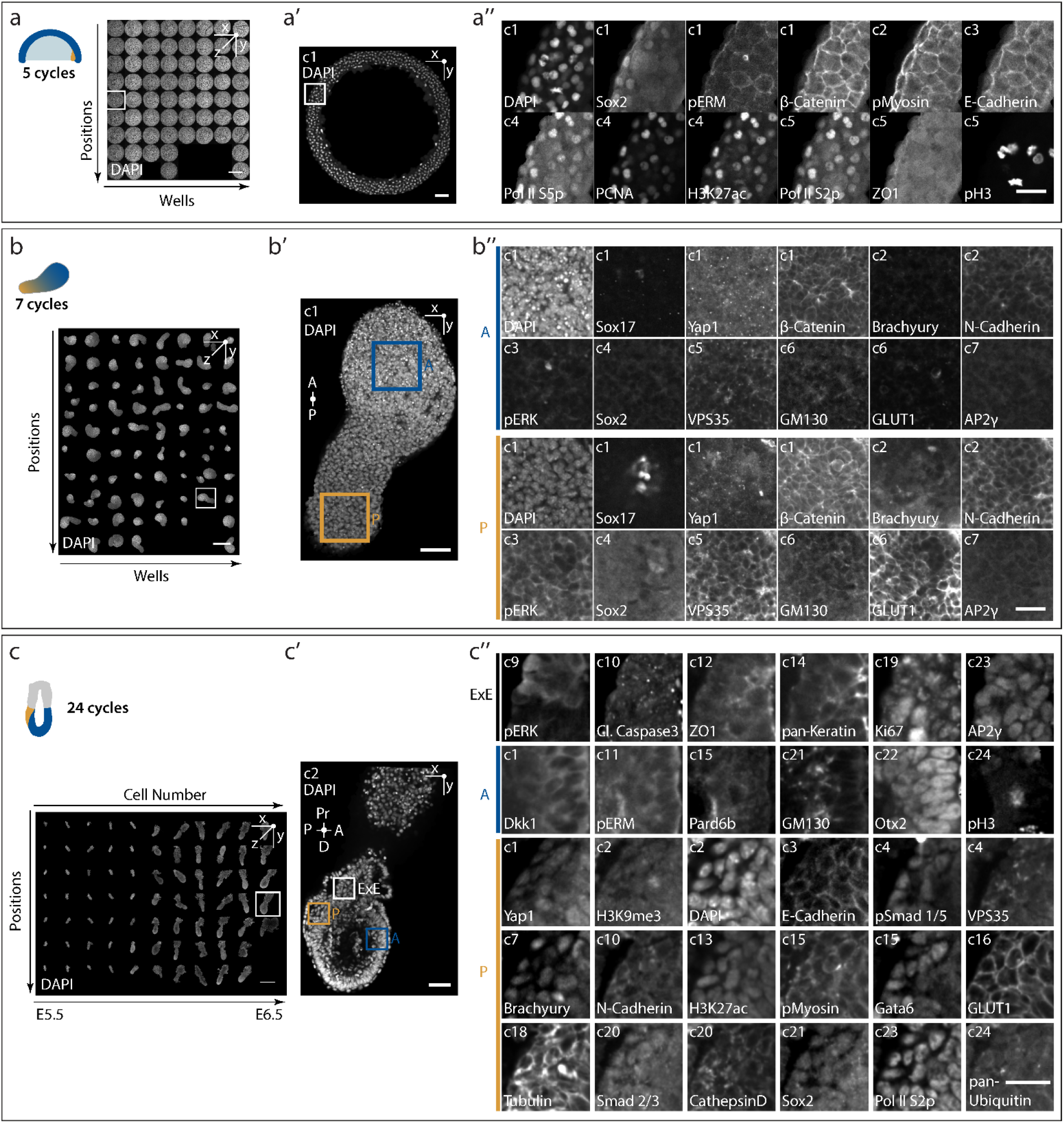
Multiplexed protein imaging of gastrulating zebrafish embryos, stem cell-derived murine gastruloids and post-implantation mouse embryos using Shield-4i. a. 3D overview of zebrafish embryos imaged in the multiplexing experiment. DAPI labels DNA. The ROI indicates the representative embryo shown in a’. Scale bar, 500 μm. a’. Representative 2D cross-section of DAPI staining in the gastrulating zebrafish embryo shown in a. The boxed region indicates the magnified inset shown in a’’. Scale bar, 50 μm. a’’. Magnified inset from a’ showing sequential Shield-4i imaging cycles detecting DAPI together with Sox2, β-Catenin, pERM, pMyosin, E-Cadherin, Pol II S5P, PCNA, H3K27ac, Pol II S2P, ZO1, and phospho-H3. Scale bar, 25 μm. b. 3D overview of murine stem cell-derived gastruloids imaged in the multiplexing experiment. DAPI labels DNA. The ROI indicates the representative gastruloid shown in b’. Scale bar, 500 μm. b’. Representative 2D cross-section of DAPI staining in the gastruloid shown in b. The blue and ochre boxed regions indicate the anterior (A) and posterior (P) magnified insets shown in b’’, respectively. Scale bar, 50 μm. b’’. Magnified insets corresponding to A (blue) and P (ochre) in b’, showing sequential Shield-4i imaging cycles detecting DAPI together with Sox17, Yap1, β-Catenin, Brachyury, N-Cadherin, pERK, Sox2, VPS35, GM130, GLUT1, and AP2γ. Scale bar, 25 μm. c. 3D overview of post-implantation mouse embryos imaged in the multiplexing experiment. DAPI labels DNA. The ROI indicates the representative embryo shown in c’. Scale bar, 640 μm. c’. Representative 2D cross-section of DAPI staining in the mouse embryo shown in c. The blue, white and ochre boxed regions indicate the anterior epiblast (A), extra-embryonic ectoderm (ExE), and posterior epiblast (P), respectively, with corresponding magnified insets shown in c’’. Pr, proximal; D, distal. Scale bar, 50 μm. c’’. Magnified insets corresponding to ExE, A and P in c’, showing sequential Shield-4i imaging cycles detecting DAPI together with pERK, cleaved caspase-3, ZO1, pan-Keratin, Ki67, AP2γ, Dkk1, pERM, Pard6b, GM130, Otx2, pH3, Yap1, H3K9me3, E-Cadherin, pSmad 1/5, VPS35, Brachyury, N-Cadherin, H3K27ac, pMyosin, Gata6, GLUT1, Tubulin, Smad 2/3, CathepsinD, Sox2, Pol II S2P and pan-Ubiquitin. Scale bar, 25 μm.

### An integrated and FAIR image analysis workflow for high-throughput 3D multiplexed image datasets

High-throughput multiplexed 3D imaging datasets easily reach tens of terabytes in size, posing substantial challenges for storage, transfer and quantitative analysis at scale. Extracting meaningful biological insights from such data therefore requires a computational framework that is scalable, reproducible, and accessible to researchers with limited scientific computing expertise. To address this need, we developed a dedicated analysis pipeline for volumetric multiplexed imaging data based on the open, cloud-native OME-Zarr file format and the recently developed Fractal platform^21^. Fractal provides an intuitive web interface for designing, executing, monitoring, and sharing modular image analysis workflows on high-performance computing infrastructure, while adhering to FAIR data principles^23^ (Fig. 3a). Together, OME-Zarr and Fractal lower the barrier to large-scale image analysis, promote reproducibility, and ensure interoperability with community bioimage viewers such as Fiji, Vizarr and napari^47–49^ (Fig. 3a).

**Fig. 3.**
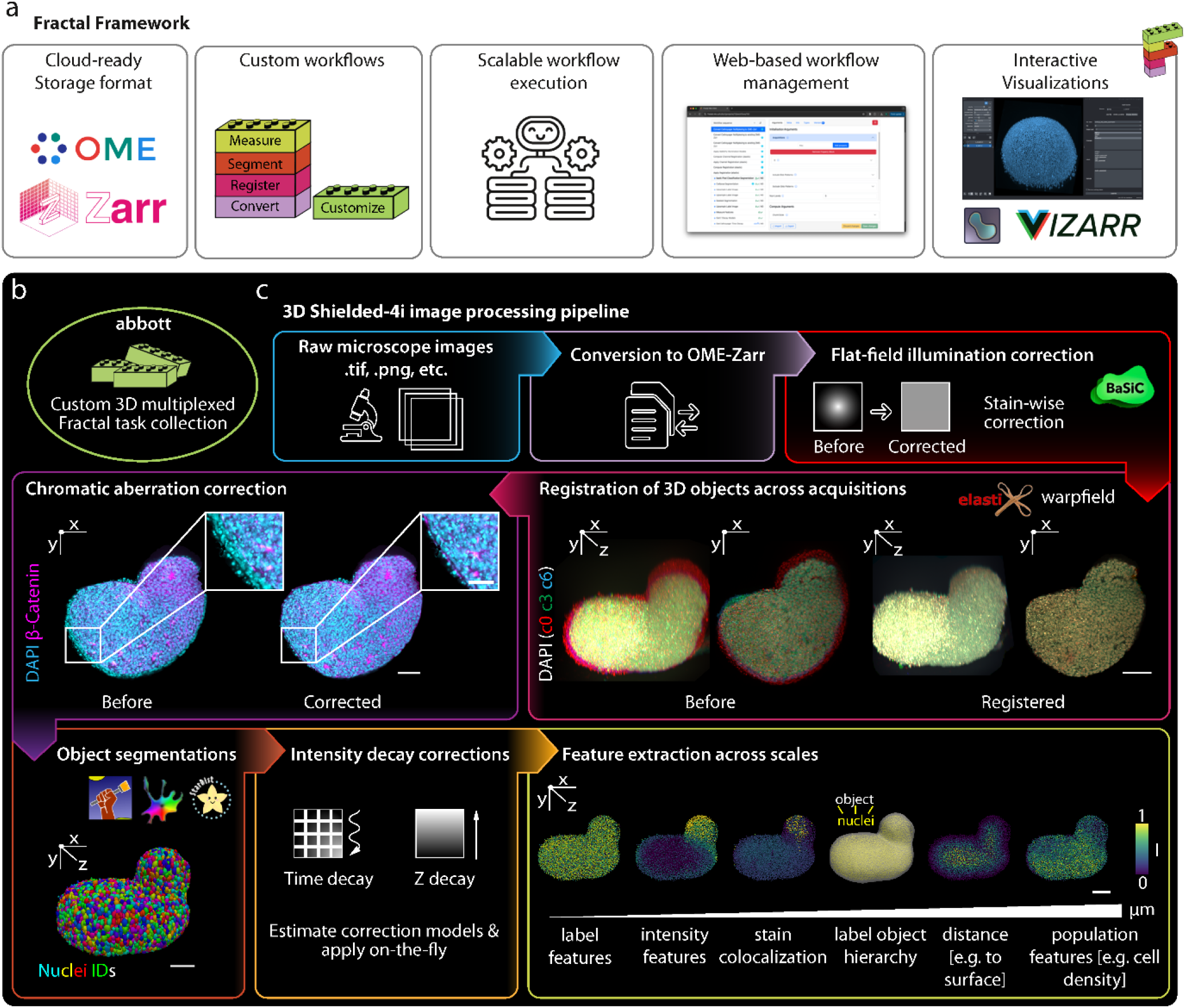
Integrated image processing pipeline for analyzing high-throughput multiplexed 3D imaging datasets. a. Overview of the Fractal framework. Fractal processes bioimage data in the OME-Zarr format, where modular and customizable workflows can be designed from reusable analysis tasks. These workflows can be executed locally or on high-performance computing clusters. Workflow design, execution, and monitoring are managed through a user-friendly web interface. Processed datasets can be interactively visualized in compatible bioimage viewers. b. The *abbott* Fractal task collection, developed in this study, extends Fractal’s capabilities for high-content 3D multiplexed imaging data. c. Standard image-processing pipeline for file conversion, illumination correction, registration of multiplexed acquisitions, chromatic aberration correction, object and single-cell segmentation, and extraction of multiscale features corrected for intensity decay along the z-axis and across imaging cycles. Individual steps are illustrated as follows: *Registration of 3D multiplexing cycles*: Overlay of DAPI signal from three imaging cycles (c0, red; c3, green; c6, blue) shown before (left) and after (right) registration for a representative gastruloid, 3D projection (left) and 2D cross-section (right). After registration, precise overlap of DAPI channels across multiplexing rounds is achieved at single-cell resolution. *Chromatic aberration correction:* Representative 2D-section showing DAPI (cyan) and β-Catenin (magenta) signals before (left) and after (right) chromatic aberration correction. *Object segmentation:* 3D projection of segmented nuclei color-coded by nuclei ID. *Intensity decay corrections:* Time and z-decay correction models are estimated per stain and applied on-the-fly during feature extraction. *Illustration of feature extraction across scales:* Feature categories extracted from multiplexed imaging data are illustrated across six panels (left to right). Label features: nuclei centroids are color-coded by nuclear roundness, derived from segmented label objects. Intensity features: nuclei are colored by the median Brachyury intensity. Colocalization features: correlation between Brachyury and Sox17 mean intensities is mapped onto nuclei centroids. Object hierarchy features: nuclei segmentations are displayed in yellow alongside the corresponding gastruloid segmentation mask in white, illustrating parent-child label relationships. Distance features: nuclei centroids are color-coded by their distance to the gastruloid surface. Population-level features: local nuclei density is represented by the number of touching neighbors per nucleus. Scale bars, 75 µm.

Building on previously published pipelines^15^, we developed three dedicated Fractal task collections, *abbott, abbott-segmentation-tasks* and *abbott-features*, to support volumetric multiplexed imaging datasets (Fig. 3b). These collections provide 3D multiplexed preprocessing modules, including file-format conversion and sample registration, object segmentation, and scale-crossing feature extraction, encompassing subcellular and cellular intensity measurements, object hierarchies, and cell-density quantification (Fig. 3c). The pipeline first converts unprocessed microscopy data into the OME-Zarr format using instrument-specific converters, many of which are available through the Fractal task library^21^. Flat-field illumination correction is then applied with BaSiCPy^50^ to compensate for uneven illumination and detection across the field of view (Methods). Whole-mount structures imaged over multiple staining cycles are registered to a reference cycle using DAPI, which labels DNA and is acquired in every cycle. To support this step, we incorporated the *elastix* and *warpfield* libraries in our task package as complementary registration strategies^24,25^. Registration proceeds through rigid, affine, and non-linear transformations, progressively increasing the degrees of freedom to correct cycle-to-cycle deformations. For multi-channel acquisitions affected by chromatic aberration, an additional channel-registration step can be applied using analogous transformations relative to DAPI (Fig. 3c and Methods). Registration quality is assessed by computing single-cell Pearson intensity correlations between each cycle and the reference acquisition (Fig. 1d), enabling poorly registered cells to be excluded from downstream analyses.

For segmentation of nuclei, cells, membranes, and whole-mount embryos or gastruloids, the pipeline supports established tools including Cellpose, Ilastik, StarDist, and PlantSeg^51–55^, allowing users to compare approaches and select the best-performing model for each tissue context (Fig. 3c). Multiscale feature extraction is then performed using the *abbot-features* library (Methods). To reduce technical intensity biases, stain-specific signal decay caused by fluorophore quenching in refractive index-matched imaging solution during acquisition, as well as z-axis attenuation caused by light scattering through tissue, are quantified and corrected^15^ (Methods). These corrections calibrate the data at single-pixel resolution across imaging rounds, channels, and tissue depth, supporting both engineered feature extraction (Fig. 3c) and data-driven learning directly from microscopy images, thereby resulting in a common quantitative representation that can be interrogated across organizational scales.

### Estimating positional information in gastruloids with multimodal measurements

Next, we combined Shield-4i with the computational pipeline developed here to assess the self-organization capacity and reproducibility of gastruloids (Fig. 4a). To quantify spatial organization, each gastruloid was fitted by a spline along its midline, defining an anterior-posterior (AP) axis and a center-surface (CS) axis perpendicular to it (Methods). Individual cell positions were then mapped into the resulting spatial reference framework (Fig. 4b-b’’). Using this approach, we analyzed the spatial profiles of protein states measured in our multiplexed dataset (Fig. 2b-b’’), reporting on lineage identity, signaling activity and cellular physicochemical states along the AP axis (Fig. 4c and Supp Fig. 3a). Consistent with previous observations^31,38,56–58^, we detected posterior enrichment of pERK, β-Catenin, Sox2 and Brachyury. In addition, the analysis uncovered AP-associated gradients that have not been previously quantified, including posterior enrichment of the metabolic marker GLUT1 and higher levels of the mechanosensor Yap1 in the anterior side. These measurements confirmed highly reproducible spatial patterns with low intra- and inter-sample variability (Fig. 4c and Supp Fig. 3a).

**Fig. 4.**
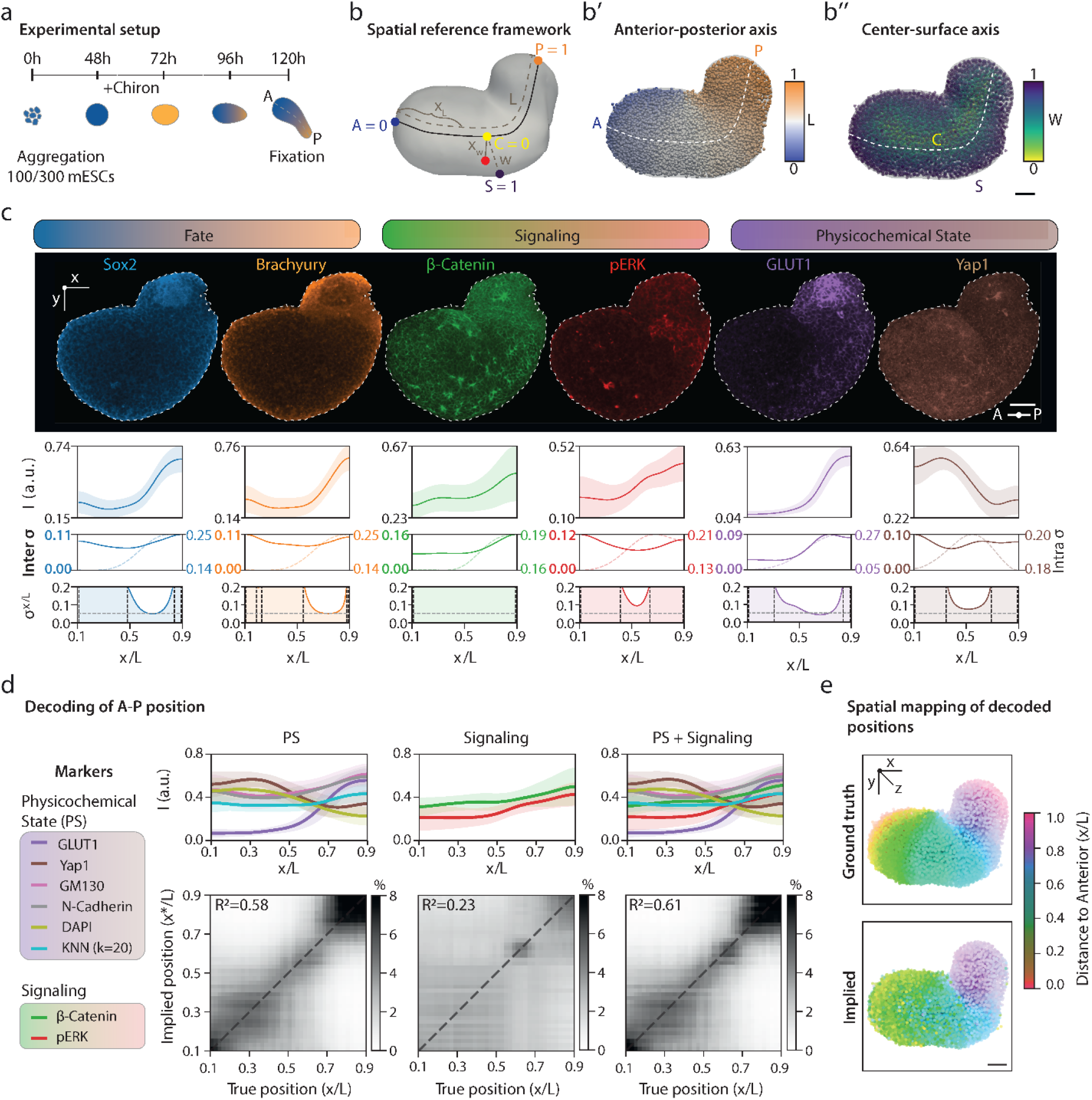
Positional information estimation in gastruloids from multimodal measurements. a. Schematic of the gastruloid formation protocol. Gastruloids were generated over 120 hours starting from 100 or 300 murine embryonic stem cells (mESCs) and pulsed with the Wnt agonist Chiron (CHIR99021) between 48 and 72 hours (Methods). b. Schematic of the spatial reference framework. A spline of length *L* (black dashed line) is fitted along the anterior-posterior (AP) axis of each gastruloid. For each nucleus, the normalized AP position is defined from A = 0 (anterior) to P = 1 (posterior) and reported as *x*/*L*; the normalized center-surface (CS) position is defined from C = 0 (center) to S = 1 (surface) and reported as *x*/*W*, where *W* is the local CS width and the variables w and w’ denote the distances used to compute the normalized CS position (Methods). b’. 3D illustration of segmented nuclei centroids of the same gastruloid as in b, color-coded by normalized AP position: blue (anterior, A = 0) to orange (posterior, P = 1). b’’. Same gastruloid as in b’, with nuclei color-coded by normalized CS position: yellow (center, C = 0) to purple (surface, S = 1). Scale bar, 75 µm (b-b’’). c. Spatial expression profiles of six multiplexed markers, grouped by functional category: Fate (Sox2, Brachyury), signaling (β-Catenin, pERK) and cellular physicochemical state (GLUT1, Yap1). *First row:* Representative 2D cross-sections of immunofluorescence signal for each marker. Scale bar, 75 µm. *Second row:* Mean intensity profiles (bold lines) along the normalized AP axis (*x*/*L*), with shaded regions indicating the standard deviation across gastruloids (n=42). *Third row*: Positional variability decomposed into inter-gastruloid variability (continuous line, left y-axis) and intra-gastruloid variability (dotted line, right y-axis). *Fourth row:* Positional error (σ_*x*/*L*_), quantifying the standard deviation of inferred positions from the expression profile of individual gastruloids plotted against *x*/*L*. Horizontal dashed line corresponds to two effective nuclei diameters, vertical dashed lines to where positional error reaches 0.2, bounding zones with low positional error. d. Decoding of AP position with an increasing number of markers in gastruloids. Top row: Mean intensity profiles (bold lines) along the normalized AP axis (*x*/*L*), with shaded regions indicating the standard deviation across gastruloids. Bottom row: Decoding maps *P*(*x*^*^|*x*) averaged across gastruloids. n=42. PS: Physicochemical State. e. Spatial mapping of ground-truth (GT, top) and decoded positions (bottom) for a single gastruloid, displayed as 3D projection. Nuclei centroids are color-coded by distance to anterior (*x*/*L*). Scale bar, 50 µm.

Positional error, estimated by relating fluctuations in single marker intensity to its local spatial gradient^59^ (Methods), was lowest near the boundary zone between anterior and posterior ends of gastruloids, where expression profiles changed most steeply (Fig. 4c and Supp Fig. 3a). To quantify the positional information encoded collectively by these multimodal measurements, we constructed a Bayesian decoder to predict individual cell positions along the AP and CS axes from signaling readouts (β-Catenin and pERK), and cellular physicochemical state features (GLUT1, Yap1, GM130, N-Cadherin, DAPI and KNN-based local density) measured here. Individual feature modules each encoded partially non-redundant positional information, and their combination enabled reconstruction of cell coordinates along the AP axis with substantial accuracy (R^2^ ∼ 0.6, Fig. 4d-e). Importantly, positional information was not distributed uniformly along the AP axis. The decoder achieved highest positional precision in the boundary zone, followed by the posterior region, between 0.7 to 0.9 normalized AP position, whereas the anterior domain, from 0.1 to 0.5 normalized AP position, exhibited substantially greater ambiguity. Prediction along the CS axis was consistently less accurate (R^2^ ∼ 0.2, Supp Fig. 3b), in line with lack of radial organization in the measured features. Together, these results demonstrate that gastruloids establish remarkably reproducible spatial patterns not only of gene expression as previously reported^59^, but also of signaling and broader cellular physicochemical states, which enable prediction of cellular position along the AP axis, albeit with region-dependent accuracy.

Finally, we asked whether the observed precision of molecular patterning is reflected in cellular decision outcomes, by quantifying the cell fate heterogeneity across gastruloids. Using a random forest classifier, cells expressing AP2γ, Brachyury, Sox17 or Sox2 were assigned to primordial germ cell-like, posterior mesodermal, endodermal or neural progenitor-like states, respectively^33,60–62^ (Supp Fig. 4a-b). We then measured variation in the abundance of each cell type and its position along the AP axis across structures. Neural and posterior mesodermal populations showed highly reproducible cell fractions between gastruloids and consistently localized to the posterior region, between 0.6 and 0.9 normalized AP position, where positional information peaked highest (Supp. Fig. 4c and Fig. 4d). In contrast, the primordial germ cell- and endodermal-like populations substantially had broader positional variability across gastruloids and showed larger differences in cell number (Supp Fig. 4c-d). These observations raise the possibility that the regional differences in positional information may contribute to the reproducibility of cell fate specification. Future studies combining multimodal measurements before and during symmetry breaking with perturbation experiments will be required to determine how positional information, molecular patterning and cell fate decisions emerge and interact to collectively drive tissue self-organization.

### Dissecting multiscale information flow in multicellular systems using Shield-4i

Cell type fractions also differed significantly between small and large gastruloids for AP2γ-, Brachyury-, and Sox17-positive cells (Supp Fig. 4c-d), pointing towards coordinated interactions between emerging fates and tissue architecture. Motivated by these observations, we next asked whether variations in cellular physicochemical state, signaling and fate specification could explain differences in gastruloid morphology. We hypothesized that molecular, cellular and tissue-scale features are statistically coupled, reflecting coordinated processes across scales during self-organization. To test this, we quantified gastruloid outlines from maximum projections using elliptic fourier descriptors^63^ (Methods) and embedded shape variation into a low-dimensional morphospace by applied principal component analysis (Fig. 5a). PC1, accounting for 83% of the observed phenotypic variability, was primarily associated with axis elongation, whereas PC2 captured variation in gastruloid thickness and accounted for a further ∼11% of the variance (Fig. 5b). We then combined the spatial molecular patterns and cell type fractions defined earlier (Fig. 4c and Supp Fig. 3a, 4), to determine whether they jointly explain gastruloid shape diversity (Fig. 5c). Remarkably, despite relying on a limited set of molecular and cellular features, the combined physicochemical, signaling and fate measurements accurately predicted the primary axis of morphological variability, PC1 (R^2^ ∼ 0.8, Fig. 5d). Feature importance analysis identified spatial patterns of pERK, Yap1, N-Cadherin and GLUT1 as strongest predictors of axis elongation (Fig. 5e-g), consistent with recent studies implicating a role for mitogenic signaling, mechanosensing, cell-cell adhesion, and metabolic state in gastruloid self-organization^39,46,57,58,64^. These findings highlight substantial shared information between molecular patterning, lineage composition, and morphological symmetry breaking, showcasing how processes operating across scales collectively shape developmental outcomes. Altogether, Shield-4i puts forward a framework for quantitatively linking signaling and cellular physicochemical state diversity to emergent tissue-level phenotypes and uncovering scale-crossing relationships in self-organizing multicellular systems.

**Fig. 5.**
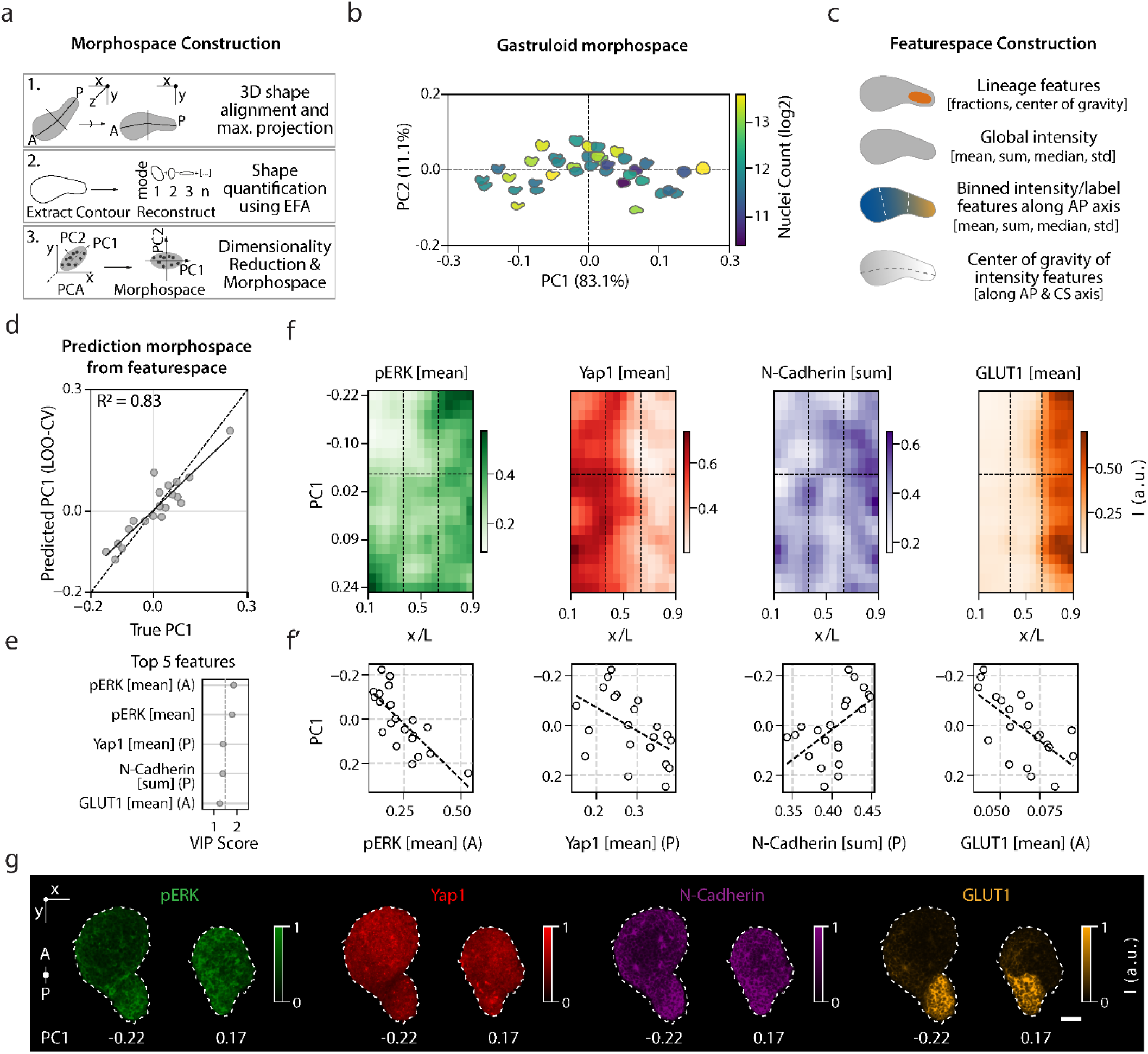
Morphospace analysis links gastruloid shape variability to molecular and lineage features. a. Workflow for morphospace construction. Gastruloids were first aligned in 3D along the anterior-posterior axis and maximum-projected to 2D. Shape outlines were quantified using elliptic Fourier analysis (EFA; Methods), followed by principal component analysis (PCA) of the resulting shape descriptors. b. Gastruloid morphospace: PCA embedding of gastruloid shape descriptors (explained-variance shown on each axis). Gastruloids are color-coded by nuclei count and displayed as their reconstructed morphologies (n=40). c. Construction of the feature space used for shape prediction: lineage features, global marker intensity features, intensity/label features binned along the AP axis, and center-of-gravity measurements for marker distributions along the AP axis (Methods). d. True versus predicted PC1 from feature-space measurements using partial least squares (PLS) regression (R^2^ = 0.83), evaluated by leave-one-out cross-validation (LOO-CV; Methods). Dashed line demarcates perfect prediction, solid line linear fit. e. Top five features associated with PC1, ranked by their Variable Importance in Projection (VIP) score (Methods). Dashed vertical line depicts VIP score ≥ 1.1. A, anterior bin; P, posterior bin. f. Kymographs of the four top features (pERK combined): gastruloids are sorted by PC1 along the y-axis, with the x-axis representing position along the anterior-posterior axis (*x*/*L*). Colors indicate the normalized intensity values of markers. Vertical dashed lines indicate anterior, center and posterior bin; horizontal dashed line, PC1 = 0. n=21. f’. Correlation between PC1 and each of the four top-ranked features. Each point is one gastruloid (n=21), dashed linear indicates linear fit. g. Representative 2D cross-sections of pERK, Yap1, N-Cadherin, and GLUT1 in two gastruloids with low PC1 (-0.22; more elongated) and high PC1 (0.17; more round). Scale bar, 50 µm.

## Discussion

Development, regeneration, and disease emerge from coordinated interactions between genetic programs, signaling networks, and physical forces. Although genetics, chemical perturbation screens, active matter physics, mathematical modeling, and machine-learning approaches have provided powerful tools for studying morphogenesis and genotype-to-phenotype relationships^65–69^, experimental frameworks that jointly capture multicellular environments, physicochemical cell states, and developmental outcomes have remained limited. Such measurements are essential for quantitatively linking molecular organization to tissue morphogenesis and for understanding how developmental information is integrated across biological scales.

Here, we introduce Shield-4i as an integrated experimental and computational framework for highly multiplexed, whole-mount protein state imaging in intact three-dimensional developmental and stem cell-derived systems. By combining SHIELD-based polyepoxide crosslinking with high-temperature SDS elution, iterative immunofluorescence imaging, and volumetric microscopy, Shield-4i enables repeated rounds of antibody staining and elution while preserving tissue morphology and antigenicity for up to 24 imaging cycles over 90 days. We demonstrate its applicability across gastrulating zebrafish embryos, post-implantation mouse embryos, and stem cell-derived murine gastruloids, achieving up to 30-plex measurements of proteins and protein states across tens of samples. In parallel, we established a scalable image analysis workflow compatible with FAIR data standards and multiscale feature extraction through the Fractal platform. Together, these advances provide a generalizable framework for building high-dimensional physicochemical cartographies of development across species and model systems.

SHIELD-mediated epoxidation preserves nucleic acids, endogenous proteins, and fluorescent signals while maintaining tissue architecture^20^. As a result, Shield-4i is readily applicable to transgenic systems in which specific proteins, signaling states, or lineages are genetically labeled. In parallel, advances in whole-mount, highly multiplexed RNA imaging have expanded the scope of spatial transcriptomics in three-dimensional tissues and embryos^70,71^. However, many of these approaches remain destructive to tissue antigenicity, limiting their compatibility with subsequent protein measurements. Given the ability of SHIELD-based protocols to preserve both nucleic acids and proteins, an exciting future direction will be the integration of highly multiplexed RNA and protein-state profiling within the same specimen.

Applying Shield-4i to gastruloids demonstrated that signaling activities, cellular physicochemical states and lineage composition establish highly reproducible spatial patterns that together encode positional information and predict tissue morphology. The strong association between molecular organization and axial elongation suggests that the physicochemical state itself acts as a continuously evolving internal representation of a cell’s developmental history, integrating prior exposure to morphogens that provide positional information^38,58,72,73^, local cellular rearrangements such as sorting and directed migration^56,74^, and changes in material properties that reinforce tissue-scale organization^75^. Extending Shield-4i to earlier stages of symmetry breaking will help define how initially subtle signaling, metabolical or mechanical biases are amplified into organized tissue-level patterns.

Although the causal relationships between these processes remain to be established, our findings suggest that information flows linking cellular physicochemical state, signaling, tissue mechanics, and lineage commitment can be traced in intact multicellular systems. Combining such measurements with temporal sampling and cell tracking^76^, aided by computational approaches that link disjointed datasets^77–79^, may enable us to study development as a system of coupled single-cell trajectories through interacting state manifolds operating across distinct timescales, where cellular decisions emerge from the progressive coordination of physicochemical state, signaling and tissue organization. This could provide a quantitative framework for understanding development and may ultimately connect developmental biology to broader theories of non-equilibrium self-organization developed in physics.

## Methods

### Sample Preparation

#### Zebrafish embryos

All animal work was carried out by the FELASA guidelines and standards of the University of Zurich. Embryos were raised at 28 °C in E3 medium until reaching between 5.5 and 6 hours post fertilization. Embryos were fixed in freshly prepared, pre-chilled 4% paraformaldehyde (PFA; Lucerna Chem) in 1X PBS. Samples were maintained on ice throughout collection and subsequently fixed overnight at 4°C on a rotary shaker for up to 18 hours. Fixation was terminated by washing the samples thoroughly five times in 1X PBS. Following fixation, embryos were manually dechorionated in a glass Petri dish under a stereomicroscope using watchmaker forceps. Permebilization was then achieved by stepwise transfer into increasing concentrations of methanol (Sigma-Aldrich), raising the methanol content in 25% increments with 5-minute incubations at each step until reaching 100% methanol. Embryos were then stored in 100% methanol at -20°C overnight. On the day of the experiment, samples were gradually rehydrated back into PBS-T (PBS supplemented with 0.1% Tween 20; Sigma-Aldrich) in 25% steps. The yolk was subsequently removed by repeated mechanical agitation in a glass centrifuge tube with a Pasteur pipette, with at least six washes performed in PBS-T.

#### Mouse embryos

Wildtype CD1 mice at 10-12 weeks of age were used in this study. The animals were maintained under a 14-hour light/10-hour dark cycle with free access to food and water. Male mice were kept individually, whereas the female mice were housed in groups of up to four per cage. Animal experiments and husbandry were performed according to the German Animal Welfare guidelines and approved by the Landesamt für Natur, Umwelt und Verbraucherschutz Nordrhein-Westfalen (State Agency for Nature, Environment and Consumer Protection of North Rhine-Westphalia, approval number 81-02.05.50.23.003). Post-implantation embryos at day 5.5 to 6.5 post fertilisation were isolated as described previously^80^. Reichert’s membrane was removed using enzymatic digestion^81^. Fixation was performed in 4% PFA for 20 minutes, followed by thorough washing of the embryos in 1X PBS.

#### Gastruloids

129/svev, EmbryoMax© CMTI-1 mouse embryonic stem cells were cultured in a humidified incubator (5% CO_2_, 37°C), and kept between passage 20 and 25. Cells were cultured in 2i/LIF medium: DMEM + GlutaMAX (Gibco) containing 10% ES-certified FBS (Gibco), non-essential amino acids (Gibco), sodium pyruvate (Gibco), beta-mercaptoethanol (Gibco), penicillin/streptomycin (Gibco), 100 ng/mL^−1^ mouse LIF (custom-produced by the EPFL Protein Production and Structure Core Facility), 3 μM GSK3 inhibitor Chiron (CHIR99021), and 1 μM MEK1/2 inhibitor (PD0325901, Selleckchem). Gastruloids were produced following a previously established protocol^27,57^. mESCs were washed with 1X PBS (Gibco), dissociated with Accutase (StemPro), centrifuged, washed, and resuspended in pre-warmed N2B27 medium comprising 50% DMEM/F12 (Gibco) and 50% Neurobasal (Gibco), supplemented with 1X N2 (Gibco) and 1X B27 (Gibco). After cell concentrations were determined using a Countess 3 automated cell counter (Invitrogen), aggregates were initiated by seeding either ∼100 or ∼300 cells per well in 40 μL of N2B27 into low-attachment, round-bottom 96-well plates (Corning), one plate per condition. At 48 h, 150 μL of N2B27 containing 3 μM Chiron was added to each well for a 24 h pulse. After the pulse, 150 μL of medium was replaced with fresh N2B27 daily. All gastruloids were fixed at 120 hours, irrespective of morphology, in freshly prepared, pre-chilled 4% paraformaldehyde in 1X PBS overnight at 4°C on a rotary shaker for up to 18 hours. Fixation was stopped by washing the samples thoroughly five times in 1X PBS.

### Mounting

Plate preparation was performed on the day of sample mounting using polystyrene flat-bottom 96-well plates (Greiner). To promote sample attachment, wells were coated with a mixture of human fibronectin (Sigma-Aldrich), diluted 1:10 in PBS, and Poly-D-Lysine (Thermo Fisher), diluted 1:2 in PBS. Each well was incubated with 50 μl of coating solution for 2 hours at room temperature, after which the solution was removed and the plate was allowed to air-dry for an additional 2 hours. Before mounting, wells were filled with 300 μl PBS. Samples were then gently transferred using a glass Pasteur pipette to minimize mechanical stress. Allowing the specimens to settle passively through the liquid column promoted a mostly reproducible orientation within the well, with the animal pole of zebrafish embryos facing the bottom of the plate, and the proximal-distal axis of mouse embryos and the anterior-posterior axis of gastruloids aligned parallel to the plate surface. Up to 10 structures were mounted per well. To reinforce adhesion, mounted samples were exposed to high-intensity 365 nm UV light for 1 hour on a UV transilluminator (Lab Gene Instruments) and imaged on a Cell Voyager 8000 microscope by acquiring a 100 μm z-stack with 4 μm spacing using the 488 nm laser at full power and 100 ms exposure. Finally, plates were stored overnight at 4 °C before further processing.

### Shield Fixation Step

Samples were subjected to an epoxy-based Shield treatment using a protocol adapted from the Chung laboratory^20^. To allow uniform reagent diffusion throughout the tissue, samples were first incubated in Shield-OFF (Lifecanvas Technologies) solution consisting of Epoxy, Shield buffer, and water, mixed at a 1:2:5 ratio, pre-cooled and maintained for 23 hours at 4 °C. Crosslinking was then initiated by replacing this solution with a pre-warmed Shield-ON solution containing Shield-ON and Epoxy at a 7:1 ratio and incubating samples for 3 hours at 37 °C, followed by a further 18-hour incubation at 37 °C in Shield-ON only solution. After crosslinking, samples were washed 10 times with 1X PBS and incubated at 37 °C for 20 hours, with additional PBS added as needed to prevent drying.

### Whole-mount Multiplexed Immunofluorescence

A unified whole-mount multiplexing immunofluorescence workflow was established by integrating conventional immunofluorescence procedures for embryos and gastruloids^57,82–84^, with Shield-based sample preservation^20^, tissue clearing, and whole-mount volumetric imaging^85^. To reduce sample loss during processing and limit well-to-well variation, all liquid-handling steps were carried out on a Bravo automated liquid-handling platform. A residual volume of 70 μl was maintained in each well throughout the protocol to prevent drying and sample disturbance during liquid exchange.

#### 1. De-lipidation or antibody elution

At each cycle, samples were subjected to an elution/de-lipidation step in 8% SDS, 100 mM sodium sulfite and 10 mM sodium borate and pH adjusted to 9.5 (in short, SDS clearing buffer) for 2 hours at 70 °C to clear the tissue and reduce residual signal from the previous staining round or lipids (in case of the first cycle).

#### 2. Permeabilization

To fully remove residual SDS, samples were subsequently washed 16 times with and incubated in 1% PBS-Triton X-100 (PBSTr) for at least 5 hours to overnight, as residual detergent interferes with antigen-antibody binding.

#### 3. Primary antibody

Primary antibodies were prepared at 2x concentration in 0.1% PBSTr and added to samples in a volume equal to the residual volume present in each well, yielding the final working concentration. Samples were then incubated overnight at 37 °C under gentle agitation on a see-saw rocker. Antibody identifiers, dilution factors and staining cycles for each model system are listed in Tables 1-3.

**Table 2:** List of primary antibodies used in the multiplexing experiment on mouse embryos, their dilutions and staining cycles.

| Cycle | Primary | Species | color channel | Clonality | Vendor | Catalogue number | Dilution [1:x ul] |
| --- | --- | --- | --- | --- | --- | --- | --- |
| 1 | Yap1 | Rabbit | red | monoclonal | Cell Signaling | 14074 | 200 |
| 1 | Dkk1 | Goat | far-red | polyclonal | R&D | AF1096 | 200 |
| 2 | H3K9me3 | Rabbit | red | polyclonal | Abcam | ab8898 | 200 |
| 3 | Oct4A | Rabbit | red | monoclonal | Cell Signaling | 83932 | 800 |
| 3 | E-Cadherin | Mouse | far-red | monoclonal | BD | 610181 | 300 |
| 4 | VPS35 | Goat | red | polyclonal | Novus | NB100-1397 | 500 |
| 4 | pSmad1/5 | Rabbit | far-red | monoclonal | Cell Signaling | 9516 | 100 |
| 7 | Brachyury | Goat | far-red | polyclonal | Thermo | PA5-46984 | 200 |
| 9 | pERK | Rabbit | green | monoclonal | Cell Signaling | 4370 | 200 |
| 10 | Cleaved Caspase-3 | Rabbit | green | monoclonal | Cell Signaling | 9664 | 400 |
| 10 | N-Cadherin | Mouse | red | monoclonal | BD | 610920 | 400 |
| 11 | pERM | Rabbit | far-red | polyclonal | Cell Signaling | 3141 | 200 |
| 12 | Sox17 | Goat | green | polyclonal | R&D | AF1924 | 400 |
| 12 | ZO1 | Rabbit | red | polyclonal | Thermo | 61-7300 | 400 |
| 13 | H3K27ac | Rabbit | far-red | polyclonal | Abcam | ab4729 | 200 |
| 14 | pan-Keratin | Mouse | red | monoclonal | Cell Signaling | 4545 | 400 |
| 15 | Pard6b | Mouse | green | monoclonal | Santa-Cruz | sc-166405 | 200 |
| 15 | Gata6 | Goat | red | polyclonal | R&D | AF1700 | 200 |
| 15 | pMyosin | Rabbit | far-red | polyclonal | Cell Signaling | 3671 | 200 |
| 16 | GLUT1 | Rabbit | red | polyclonal | Sigma | 07-1401 | 500 |
| 18 | Tubulin | Rat | far-red | monoclonal | Merck | MAB1864 | 4000 |
| 19 | Ki67 | Rabbit | far-red | monoclonal | Cell Signaling | 9129 | 200 |
| 20 | Cathepsin D | Goat | red | polyclonal | Labforce | AB0043-200 | 250 |
| 20 | Smad 2/3 | Rabbit | far-red | monoclonal | Cell Signaling | 8685 | 200 |
| 21 | Sox2 | Rabbit | green | polyclonal | Abcam | ab97959 | 200 |
| 21 | GM130 | Mouse | red | monoclonal | BD | 610823 | 200 |
| 22 | Otx2 | Goat | far-red | polyclonal | R&D | AF1979 | 400 |
| 23 | AP2γ | Mouse | green | monoclonal | Santa-Cruz | sc-12762 | 50 |
| 23 | Pol II S2p | Rat | far-red | monoclonal | Sigma | 04-1571 | 400 |
| 24 | Phospho-H3 | Rabbit | green | polyclonal | Cell Signaling | 9701 | 200 |
| 24 | pan-Ubiquitin | Mouse | red | monoclonal | Enzo | BML-PW8810-0100 | 200 |

**Table 3:**
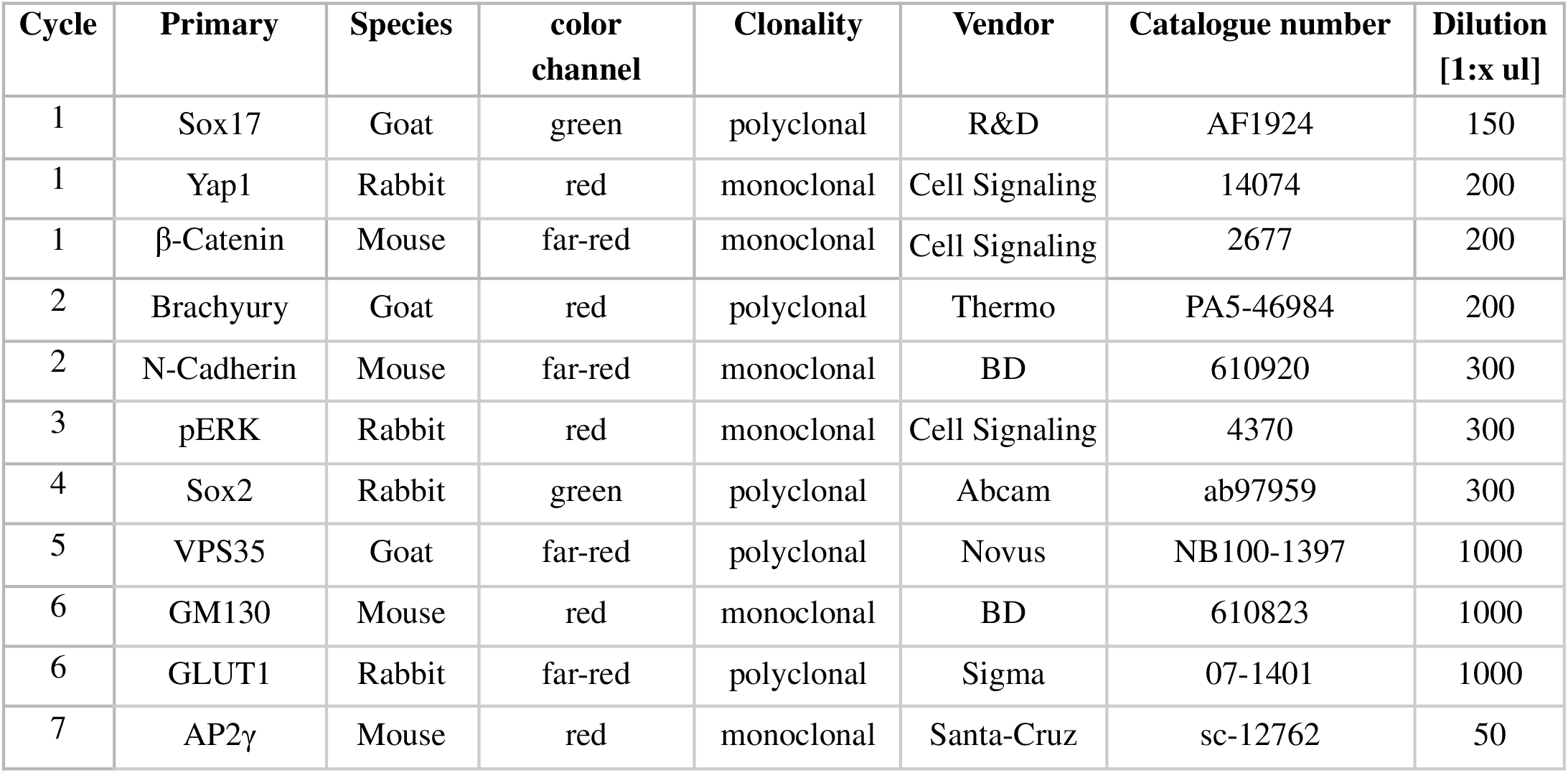
List of primary antibodies used in the multiplexing experiment on gastruloids, their dilutions and staining cycles.

| Cycle | Primary | Species | color channel | Clonality | Vendor | Catalogue number | Dilution [1:x ul] |
| --- | --- | --- | --- | --- | --- | --- | --- |
| 1 | Sox17 | Goat | green | polyclonal | R&D | AF1924 | 150 |
| 1 | Yap1 | Rabbit | red | monoclonal | Cell Signaling | 14074 | 200 |
| 1 | β-Catenin | Mouse | far-red | monoclonal | Cell Signaling | 2677 | 200 |
| 2 | Brachyury | Goat | red | polyclonal | Thermo | PA5-46984 | 200 |
| 2 | N-Cadherin | Mouse | far-red | monoclonal | BD | 610920 | 300 |
| 3 | pERK | Rabbit | red | monoclonal | Cell Signaling | 4370 | 300 |
| 4 | Sox2 | Rabbit | green | polyclonal | Abcam | ab97959 | 300 |
| 5 | VPS35 | Goat | far-red | polyclonal | Novus | NB100-1397 | 1000 |
| 6 | GM130 | Mouse | red | monoclonal | BD | 610823 | 1000 |
| 6 | GLUT1 | Rabbit | far-red | polyclonal | Sigma | 07-1401 | 1000 |
| 7 | AP2γ | Mouse | red | monoclonal | Santa-Cruz | sc-12762 | 50 |

#### 4. Secondary antibody

Following primary antibody incubation, unbound antibodies were removed by extensive washing in 0.1% PBSTr.

Secondary antibodies were prepared at 2x concentration (1:200) in 0.1% PBSTr and supplemented with DAPI at a dilution of 1:20 (2x, Invitrogen). The solution was then added in a volume equal to the residual volume in each well and incubated with the samples for 3 hours at 37 °C under gentle agitation in the dark. Details of the secondary antibodies are provided in Table 4.

**Table 4:**
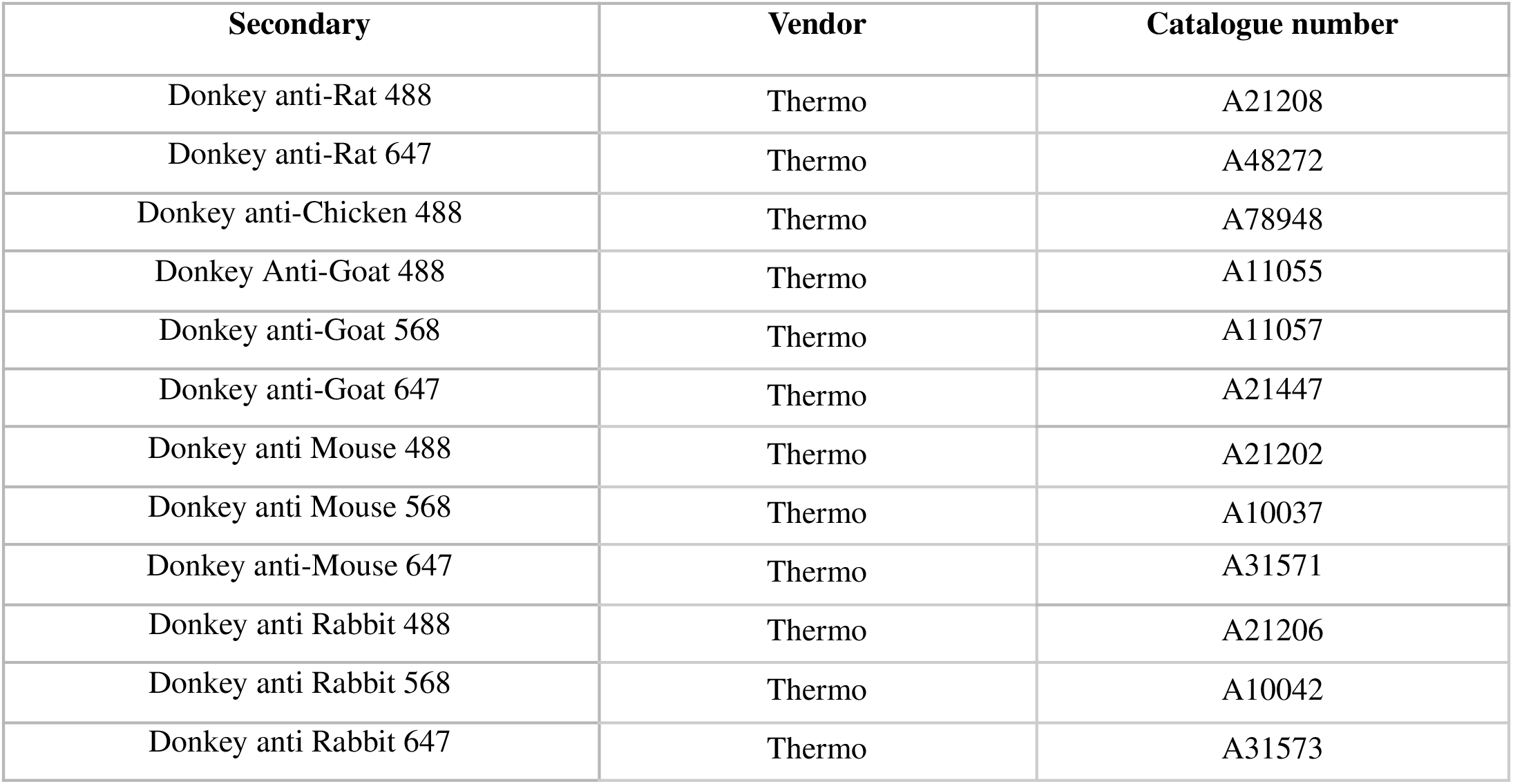
List of secondary antibodies used in the multiplexing experiments.

#### 5. Refractive index matching

After extensive washing in 0.1% PBSTr to remove excess secondary antibodies and dyes, samples were transferred to the refractive index matching solution EasyIndex (LifeCanvas Technologies). DAPI was included in the imaging solution at the same concentration used during staining to minimize its gradual loss from the sample during prolonged image acquisition. Samples were incubated in EasyIndex for at least 30 minutes before imaging to allow equilibration.

#### 6. Imaging

Images were acquired on a Yokogawa Cellvoyager 8000 spinning-disk inverted confocal microscope equipped with a 50 μm pinhole disk and a Hamamatsu ORCA Flash 4.0 camera. High-resolution volumetric imaging was performed using a 20×/1.0 NA water-immersion objective, with one or two structures imaged per field of view. Structures were detected and selected using a custom Python script. For each sample, a z-stack spanning 300-440 μm was acquired with a step size of 1 μm, depending on the thickness of structures. Laser power and exposure time were optimized individually for each stain. Imaging was performed overnight.

#### 7. Washout of refractive index matching solution

Following image acquisition, EasyIndex was removed by repeated washes and incubated in 0.1% PBSTr for at least 1.5 hours at 37 °C before starting the next staining cycle.

### Antibody Panels

#### Elution Controls

To verify efficient antibody removal between cycles, elution controls were performed by omitting primary antibodies in the subsequent round and incubating samples with secondary antibodies only.

#### File Conversion

Raw microscopy TIFF files were converted to OME-Zarr format using the *Convert Cellvoyager Multiplexing to OME-Zarr* task in *fractal-tasks-core* (https://github.com/fractal-analytics-platform/fractal-tasks-core/tree/1.5.5). For multiplexed datasets, additional imaging cycles were appended to existing OME-Zarr files using the *Convert Cellvoyager Multiplexing to Existing OME-Zarr* task in *abbott* (https://github.com/pelkmanslab/abbott). Each well in the multiwell plate was stored as a single tiled OME-Zarr image per cycle, with all fields of view stitched into a unified array based on stage-position metadata. ROI tables recording the original field-of-view coordinates were retained alongside each image to enable field-of-view-level processing in downstream analyses.

#### Illumination Bias Correction

To correct for uneven microscope illumination, flatfield corrections were estimated as described previously^15,50^. In brief, 7-diethylamino 4-methyl coumarin (Sigma-Aldrich, 0.5 mg/ml), Fluorescein (Sigma-Aldrich, 1 mg/ml), Rose Bengal (Sigma-Aldrich, 1 mg/ml) and acid blue 9 (TCI Chemicals, 10 mg/ml) were diluted in DMSO and added to a Greiner 96-well plate. For each wavelength, multiple imaging stacks were acquired across randomly selected wells. Flatfield corrections were estimated within the Fractal framework using *APx-fractal-task-collection*’s *Calculate BaSiCPy Illumination Models* task on 125 maximum projected imaging stacks per wavelength (https://github.com/pelkmanslab/APx_fractal_task_collection) and applied using *Apply BaSiCPy Illumination Models* task from the same collection.

#### Image Registration

To correct for sample shifts and deformations across imaging cycles, arising from buffer exchanges and microscope stage drift, cycle-to-cycle image registration was performed using the DAPI stain, acquired in every cycle, as a common reference. Registration transforms were estimated as a sequence of three increasingly flexible transformations (rigid, affine, and b-spline) to achieve high-quality alignment at single-cell resolution. The specific registration strategy varied by sample type: zebrafish embryos and gastruloids were registered using the elastix library throughout; for mouse embryos, rigid and affine (linear) registrations were performed using the *elastix* library, while b-spline (non-linear) deformable registration was performed using the *warpfield* library. Cells with poor registration quality (Pearson’s correlation coefficient of DAPI signal across cycles < 0.8) were automatically flagged and excluded from downstream analysis.

Channel registration was performed for datasets affected by chromatic aberration (multiplexed zebrafish and gastruloid datasets) to align all channels to the DAPI reference. Where sufficient inter-channel similarity existed, as was the case for the multiplexed zebrafish dataset, a similarity transformation was estimated directly from stacked image intensities across channels to align using the *Compute Channel Registration (elastix)* task in *abbott*. In datasets with lower inter-channel similarity, a foreground mask was first generated per channel using a dedicated Ilastik pixel classification model via the *Ilastik Pixel Classification Segmentation* task of *fractal-ilastik-tasks (https://github.com/fractal-analytics-platform/fractal-ilastik-tasks)* and the resulting embryo masks were used to estimate a per-ROI and per-acquisition affine transformation between the DAPI channel and the combined remaining channels using the *Compute Channel Registration (elastix)* task in *abbott*. In both cases, the estimated transformations were applied to the image data using the *Apply Channel Registration (elastix)* task of *abbott*.

#### Object Segmentations

Nuclei and cell segmentation in zebrafish embryos was performed on DAPI and E-Cadherin staining, respectively. Nuclei were segmented using the Cellpose library with the pre-trained *nuclei* model (diameter = 25) via the *Cellpose Segmentation* task implemented in *fractal-tasks-core*^*21,52*^. Cell segmentation was subsequently performed using a seeded watershed approach, in which nuclei labels served as seeds and E-Cadherin staining defined cell boundaries, implemented via the *Seeded Watershed Segmentation* task in *abbott-segmentation-tasks* (https://github.com/fractal-analytics-platform/fractal-cellpose-sam-task). To restrict signal quantification to the cell membrane, cell segmentation masks were eroded to a few-pixel wide membrane label using the *Reduce Cell to Membrane Segmentation* task in *abbott-segmentation-tasks*..

For mouse embryos, nuclei and cells were segmented in 3D using Cellpose-SAM applied to DAPI and E-Cadherin staining, respectively, with default settings, via the *Cellpose SAM Segmentation* task in *fractal-cellpose-sam-task* (https://github.com/fractal-analytics-platform/fractal-cellpose-sam-task). For gastruloids, nuclear segmentation was performed on DAPI staining using the *Stardist Segmentation* task within the *abbott-segmentation-tasks* collection, employing a previously custom trained 3D StarDist model^54,86^. In all cases, segmentation outputs were filtered by excluding objects falling outside a predefined size range to remove segmentation artefacts.

Whole-embryo and gastruloid segmentation was performed using a dual-channel Ilastik pixel classification model trained on nuclear (DAPI) and membrane (E-Cadherin/β-Catenin) staining from the reference cycle. Pixel classification and thresholding were carried out using the *Ilastik Pixel Classification Segmentation* task of *fractal-ilastik-tasks* with a probability threshold of > 0.8 and a minimum object size of 100,000 voxels.

### Feature Extraction and Aggregation

Feature extraction was performed as described in^15^, using the *Measure Feature Task* from the *abbott-features* task-collection (https://github.com/pelkmanslab/abbott-features). Briefly, the following feature types were extracted based on the ITK library^87^]:

Hierarchical features establish child-parent relationships across object hierarchies (e.g., linking each nucleus to its corresponding cell label), enabling feature extraction to be propagated consistently across segmentation levels.

Label features are derived from label images (e.g., nuclei, cell segmentations) and include the label identifier, centroid, orientation, bounding box, and morphological descriptors such as eccentricity, roundness, volume, etc.

Intensity features quantify the intensity distribution within each labelled region using statistical metrics including mean, median, sum and kurtosis.

Colocalization features measure the correlation or mutual information between intensity distributions across labelled regions, which is useful for quantifying protein co-occurrence or estimating image registration quality.

Distance features characterize the spatial relationship between a labelled child region and a reference parent object (e.g., the distance from a nucleus to the tissue surface). These are computed via Euclidean distance transforms and summarized using statistical metrics such as minimum, maximum, and median distance.

Population features describe the spatial context of labelled regions through graph-based representations in which objects serve as nodes and edges encode spatial relationships. Four neighborhood definitions are supported: Radial neighborhood: objects within a defined radius are considered neighbors. K-nearest neighbors (KNN): each object is linked to its *k* closest neighbors, regardless of absolute distance. Touch neighborhoods: neighbors are defined by shared surface area between objects. Delaunay triangulation (DT): neighbors are determined by applying Delaunay triangulation to object centroids.

Feature tables across OME-Zarr images were subsequently merged into a single consolidated feature table using the *Aggregate Feature Tables* task of *abbott-features*.

### Intensity Decay Corrections

Prior to final feature extraction, intensity decay along both the temporal and axial (z) dimensions was corrected on a per-stain basis using mean uncorrected intensity features. Extended acquisition times cause progressive fluorophore quenching in the refractive-index matched solution, introducing a time-dependent intensity bias if left uncorrected. For the multiplexed gastruloids, a linear model was fitted per stain to account for this temporal decay, determined using the *Get Cellvoyager Time Decay* task of *abbott-features*.

Along the z-axis, light scattering and refractive index mismatches attenuate fluorescence signal with imaging depth. Given the large number of samples mounted at varying rotations, axial intensity decay was assumed to be of technical rather than biological origin. Two correction models are available: a *1D z-model*, which fits a single decay function along the entire light path, and a *2D z-model*, which partitions the light path into medium and sample components to account for their distinct scattering properties. For the gastruloid dataset, an additional axial decay was corrected using a single-exponential 1D model, implemented via the *Get Z Decay Models* task of *abbott-features*. All intensity decay corrections were applied to the images on-the-fly during final feature extraction.

### Gastruloid Radial Intensity Decay Correction

To correct for radial intensity decay caused by high-affinity antibodies and thick gastruloids, a linear correction was fitted independently for each of the 12 multiplexed markers. Nuclei were stratified into 10 equally spaced bins along the normalised AP axis, within which intensity outliers were clipped to the 5-95th percentile range, followed by min-max normalisation. This binning decouples the radial intensity correction from underlying AP expression gradients. The normalised intensities were then binned into 5 equally spaced intervals along the normalised center-surface (CS) axis and the median intensity per interval was computed. A linear model was fitted to these five binned medians via non-linear least-squares optimisation, yielding per-marker radial decay slopes and intercepts. The resulting correction factors were stored per CS bin and marker for downstream radial decay correction.

### Spline Fitting and Reference Coordinate System

Splines were fitted to the anterior-posterior (AP) axis of each gastruloid, similar to previous approaches^88^. Briefly, segmentation masks were first smoothed, and both the mask and the Brachyury fluorescence channel were resampled to isotropic resolution. Foreground voxel coordinates were then extracted from the segmentation mask and assembled into a point cloud. An elastic principal curve was fitted to this point cloud using the ElPiGraph algorithm (NumNodes scaled dynamically with gastruloid length, λ = 0.05, μ = 0.6)^89^. Node positions and edge connectivity were rasterized into a binary skeleton by drawing line segments between connected node pairs, which was refined by morphological skeletonization to yield a single-voxel-wide centerline. The centerline was converted to a graph using the SkelePlex library^90^ and extended to the segmentation boundary by extrapolating the tangent vector at each endpoint. The posterior pole was assigned to the endpoint closest to the center of mass of the Brachyury signal.

To embed each cell in a unified reference coordinate system, two positional coordinates were computed per cell: an AP position and a radial center-surface (CS) position. The AP position was approximated as the normalized parameter value of the closest of 100 uniformly sampled points along the gastruloid spline to the cell centroid. For the radial coordinate, the gastruloid surface was first reconstructed by applying the marching cubes algorithm to the gastruloid label mask (implemented in scikit-image^91^). The normalized radial distance was then defined as: 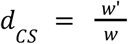 where w’ is the distance from the nucleus centroid x to the center (C) along the AP spline, and w is the distance from C to the the nearest point on the gastruloid surface, such that 0 indicates a cell on the AP axis spline and 1 a cell at the surface.

### Positional Error Estimation Along Anterior-Posterior Axis

To characterize the spatial distribution of markers along the AP axis of gastruloids, intensity profiles were computed for each gastruloid as a function of distance to the anterior pole (*x*) normalized to the gastruloid length (L). Nuclei at the anterior or posterior extremes (*x*/*L* < 0.1 or *x*/*L* > 0.9) were discarded to avoid edge artefacts. Radial intensity correction was applied per gastruloid as described above and intensities were clipped at 5th and 95th percentile and min-max normalised. Nuclei-level measurements were interpolated onto a common AP grid of 60 equally spaced points, spanning the observed *x*/*L* range. Small measurement gaps were filled by linear interpolation and the resulting profile was smoothed with a Gaussian kernel (σ = 5 grid points). Inter-gastruloid variability was quantified as the standard deviation across the smoothed per-gastruloid mean profiles. Intra-gastruloid variability was estimated at each grid point as a Gaussian-weighted local variance: for each grid point, a kernel with a bandwidth equal to three times the grid spacing was applied to the nuclei-level measurements within that bin, from which the standard deviation was derived. Both inter-and intra-gastruloid variance profiles were additionally smoothed with the same Gaussian filter.

Positional error (σ_*x*/*L*_), that is the uncertainty in inferring a cell’s position from its measured intensity, was estimated following the framework of Merle et al. (2024)^59^. For each position along the AP-grid, the positional error was defined as: 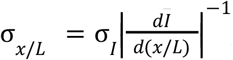, where σ_*I*_ is the inter-gastruloid standard deviation of intensity and 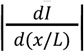 is the absolute value of the local slope of the mean expression profile 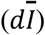. The derivative was estimated at each grid point by fitting a third-order polynomial to a sliding window of 15 consecutive grid points, taking the linear coefficient of the fit evaluated at the window center as the local slope. Positional error values exceeding the 95th percentile were clipped to suppress numerical artifacts arising at regions where the mean profile is locally flat.

### Positional Prediction

To test whether single-nucleus multimodal intensity and neighborhood measurements encode positional information within gastruloids, a Bayesian decoder was constructed to infer AP and CS position, similar to previous work^92^. Nuclei within the outer 10% of either axis were excluded to minimise edge artefacts. Prior to training, intensity measurements were corrected for radial signal decay as described above, clipped to the 1st-99th percentile and min-max normalised per gastruloid. Each axis was discretised into equally spaced bins (AP: 40 bins, ∼401 µm mean gastruloid length; CS: 8 bins, ∼85 µm mean gastruloid radius).

Separate decoders were constructed for the AP and CS axes using the same set of features (f). Feature number was first reduced by greedy permutation importance analysis prior to decoding. The reduced panel used for positional prediction consisted of the following eight features grouped into signaling (pERK and β-Catenin) and physicochemical state (GLUT1, Yap1, GM130, N-Cadherin, DAPI and KNN-density (k = 20)) modules. To compare the positional information contained in different feature classes, decoders were trained using signaling features, physicochemical state features, or the combined reduced feature panel.

For each axis, Bayes’ rule was used to infer the posterior probability that a nucleus with measured feature values ({*f*_*i*_}) originated from positional bin (*x*^*^):

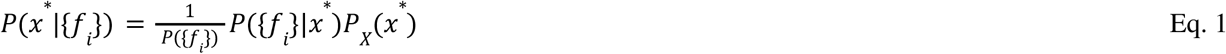

Here, *P*({*f*_*i*_}|*x*^*^) denotes the multivariate Gaussian likelihood of observing the feature vector ({*f*_*i*_}) at position (*x*^*^), *P*_*X*_ (*x*^*^) is the prior probability of a nucleus occupying position (*x*^*^) and *P*({*f*_*i*_}) is a normalization factor. A uniform positional prior was used. The parameters of the multivariate Gaussian likelihood were estimated globally across gastruloids by grouping nuclei according to their positional bin. For each gastruloid α, a decoding map was generated per each axis by inserting the measured expression levels 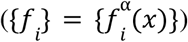 at actual position *x* into the posterior:

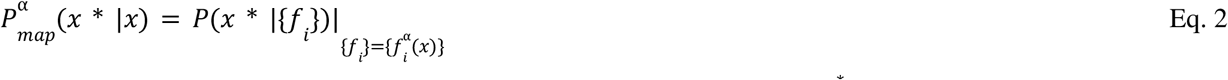

These maps represent the probability distribution of implied positions *x*^*^ for nuclei located at actual position *x*. Decoding maps were estimated separately for each gastruloid, but plotted as an average map over all gastruloids.

### Cell Type Classifier

For cell type annotation of gastruloids, multichannel intensity images containing fate markers alongside single-cell segmentation masks (e.g., nuclei) with associated feature measurements were loaded into the *napari* viewer^48^. Cell type identities were interactively annotated using the *napari feature classifier* plugin, which trains a random forest classifier on user-defined labels to generate predictions across all segmented objects^21^ (https://napari-hub.org/plugins/napari-feature-classifier.html). This workflow was applied to classify the following cell type classes: primordial germ cell-like (Ap2γ+), posterior mesodermal (Brachyury+), endodermal (Sox17+) and neural progenitor-like states (Sox2+). Gastruloids were binned into small or large categories based on total nuclei number, according to a user-defined threshold.

### Gastruloid Morphospace Construction

#### 3D shape alignment

Gastruloid label masks were resampled to isotropic resolution and the rotation of each gastruloid was normalised by aligning the AP skeleton to the x-axis via Singular Value Decomposition (SVD) and Rodrigues’ rotation formula. The widest transverse axis, used to determine the projection direction, was identified by projecting the rotated skeleton coordinates onto the zy-plane and applying a second SVD. A single affine transform containing the two rotations was then applied to the gastruloid. Anterior-posterior orientation was normalised by comparing the gastruloid and spline directionality, applying a reflection along the x-axis if needed. Maximum projection along the z-axis was then computed and the label contour was extracted (implemented in scikit-image^91^). Dorsoventral orientation was enforced along the y-axis by comparing the y-centroid of the 2D projected label with the y-centroid of its contour; as contour centroid is displaced away from any concavity, this comparison indicates the concavity direction, allowing to consistently orient the concave side downwards.

#### Shape representation

Elliptic Fourier Descriptors (EFDs, order 10) were computed on the aligned and oriented contours using the pyefd library, yielding a 40-dimensional coefficient vector per gastruloid^93^ (https://github.com/hbldh/pyefd). Coefficients were normalised by the first harmonic’s a_1_ coefficient to remove size dependence from the shape quantification. Each contour was additionally reconstructed, rescaled to unit span, and EFDs re-fitted to fully decouple shape from size and position.

#### Morphospace

Principal component analysis (PCA) was performed on the normalised EFD matrix after removing shape outliers by Mahalanobis distance in a two-dimensional space.

### Feature Space Construction for Morphology Predictions

Lineage Features were computed as the fraction of cells of each annotated cell type relative to the total cell count per gastruloid, along with the normalised center of gravity along the AP and CS axes.

Global Intensity Features: For each 3D gastruloid, intensity summary statistics (mean, sum, median, and standard deviation) were computed across all cells.

Center of Gravity of Intensity Features: For each 3D gastruloid, the normalised center of gravity of per-cell intensity values was estimated along the normalised AP and CS axis.

Binned Intensity and Label Features: Each gastruloid was divided into anterior, mid and posterior segments along a fitted AP axis spline. Intensity and label features were normalised per gastruloid and computed per segment, summarised as the mean, sum, median and standard deviation per segment.

### Prediction Morphospace From Feature Space

To test whether features spanning multiple organisational scales (see Feature space Construction) can collectively predict gastruloid morphology, we applied a two-stage regression framework to the primary axis of morphological variation (PC1, 83% explained variation). Features were standardised prior to modelling. LASSO regression was used to identify a sparse subset of informative predictors across 60 logarithmically spaced regularisation values (α = 10^−4^ to 10^0^), which were then passed to Partial Least Squares (PLS) regression with up to 2 components (both implemented in scikit-learn^91^). The optimal regularisation strength and number of components were selected by maximising leave-one-out cross-validation (LOO-CV) R^2^. To interpret the final model, feature importance was assessed using Variable Importance in Projection (VIP) scores. Features with VIP ≥ 1.1 were considered important.

## Code Availability

The code to reproduce all figures, as well as Fractal workflow files that include a comprehensive definition of the order of tasks used, their versions and all their parameters that were used to run the 3D-multiplexed image pre-processing pipeline is available in the dedicated Github repository https://github.com/pelkmanslab/shield4i_multiplexing_paper_2026.

## Data Availability

The multiplexed gastruloid and zebrafish datasets will be made available at the time of publication. The multiplexed mouse embryo time course dataset is available upon reasonable request.

## Acknowledgements

We thank members of the Pelkmans, Gilmour, Oates, and Bedzhov groups, as well as the Center for Microscopy and Image Analysis and the BioVisionCenter for helpful discussions. We are also thankful to Alexandre Mayran, for their support in establishing the gastruloid protocol in the Oates Lab and feedback on the manuscript. R.H. was supported by University of Zurich Candoc and Promoter Foundation Grant. S.S. was supported by postdoctoral fellowships from EMBO (ALTF 851-2022) and the SNSF (TMPFP3_217219), as well as an SNSF Ambizione grant (PZ00P3_216331). F.N.A was supported by a postdoctoral fellowship from the Peter and Traudl Engelhorn Foundation. D.G. was supported by the Swiss National Science Foundation (SNSF; grant 310030_204834). L.P. was supported by the SNSF (grant 310030_192622), the European Research Council Advanced Grant CROSSINGSCALES (885579), and the University of Zurich.

## Author Contributions

R.H., S.S., and L.P. designed the research. R.H. and S.S. performed the multiplexing experiments and developed the computational pipeline. F.N.A. generated gastruloids. R.F. prepared mouse embryo samples under the supervision of I.B. M.H., F.C, J.L., and V.U. contributed to image and data analysis and development of the Fractal platform. A.C.O., I.B., D.G., and V.U. provided conceptual input and project support. S.S., R.H., and L.P. wrote the manuscript with input from all authors.

## Supplementary Figures

**supplementary Fig. 1.**
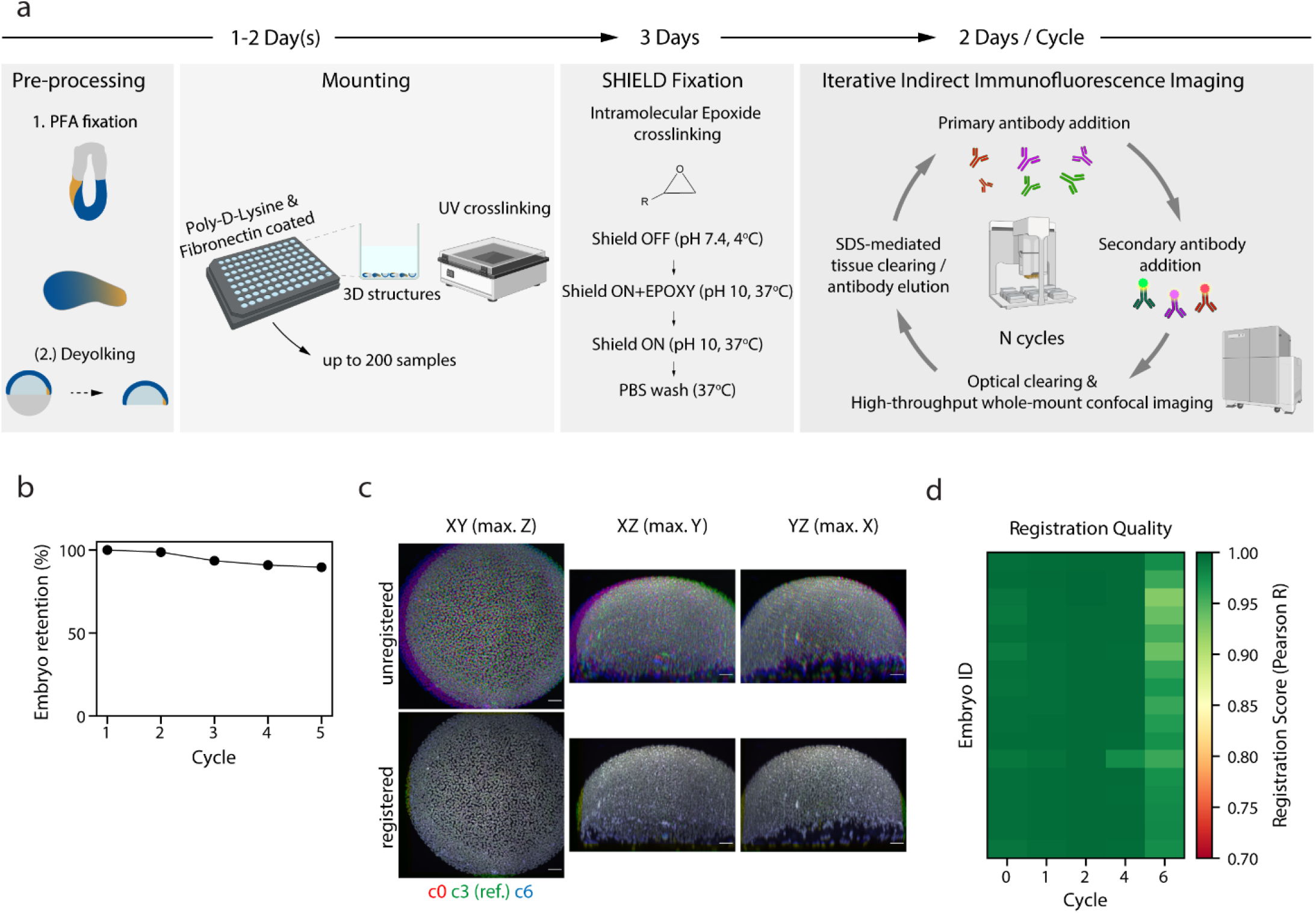
Shield-4i enables repeated imaging and accurate registration across acquisition cycles. a. Schematic overview of the Shield-4i workflow applied to gastrulating zebrafish embryos, mouse embryos, and murine stem cell-derived 3D gastruloids. PFA-fixed samples are first processed in a sample-specific manner, including deyolking for zebrafish embryos. Pre-processed samples are immobilized in Poly-D-Lysine/fibronectin-coated wells and laser-crosslinked to promote stable attachment. Samples then undergo epoxy-based Shield fixation using Shield-OFF and Shield-ON buffers to preserve tissue integrity during subsequent processing (Methods). Shield samples are subjected to repeated cycles of SDS-based elution/clearing and immunofluorescence staining, enabling up to N rounds of multiplexed 3D protein imaging. Further details are provided in Methods. b. Retention rate of mounted zebrafish embryos over five sequential acquisition cycles. c. Representative DAPI images of an exemplary zebrafish embryo acquired across all cycles, shown before registration (top) and after registration (bottom). The first cycle (c0) is shown in red, the reference cycle (c3) in green, and the final cycle (c6) in blue; improved overlap after registration indicates accurate alignment across cycles. Scale bars, 50 µm. d. Quantification of registration quality using the Pearson correlation coefficient of nuclear DAPI signal between each acquisition cycle and the reference cycle 3.

**supplementary Fig. 2.**
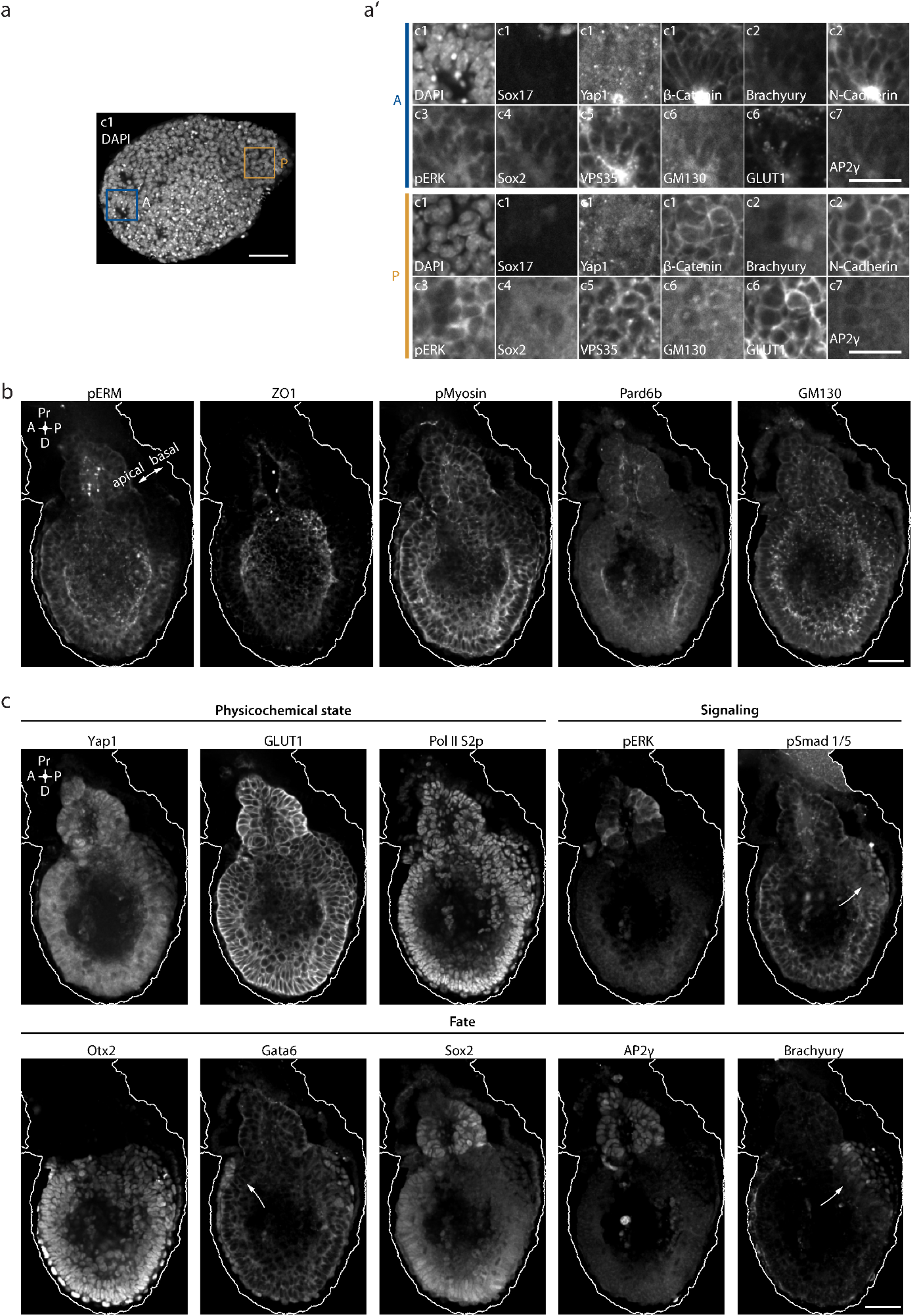
Multiplexed protein imaging of stem cell-derived murine gastruloids and characterisation of apico-basal polarity, cellular physicochemical state, signaling and cell fate markers in E6.5 mouse embryos. a. Representative 2D cross-section of DAPI staining in a gastruloid formed from 100 mouse embryonic stem cells. The blue and ochre boxed regions indicate the anterior (A) and posterior (P) magnified insets shown in a’, respectively. Scale bar, 50 μm. a’. Magnified insets corresponding to A (blue) and P (ochre) in a, showing sequential Shield-4i imaging cycles detecting DAPI together with Sox17, Yap1, β-Catenin, Brachyury, N-Cadherin, pERK, Sox2, VPS35, GM130, GLUT1, and AP2γ. Scale bar, 25 μm. b. 2D cross-sections of E6.5 mouse embryo stained for markers of apico-basal polarity (pERM, ZO1 and Pard6b) as well as markers enriched at the apical end (pMyosin and GM130). The continuous line marks the basal surface of the embryo. Pr, proximal; D, distal. Scale bar, 50 μm. c. 2D cross-sections of an embryo shown for exemplary cellular physicochemical state (Yap1, GLUT1 and Pol II S2p), signaling activity (pERK, pSmad 1/5) and cell fate markers(Otx2, Gata6, Sox2, AP2γ and Brachyury). Arrows highlight marker localisation. Scale bar, 50 μm.

**supplementary Fig. 3.**
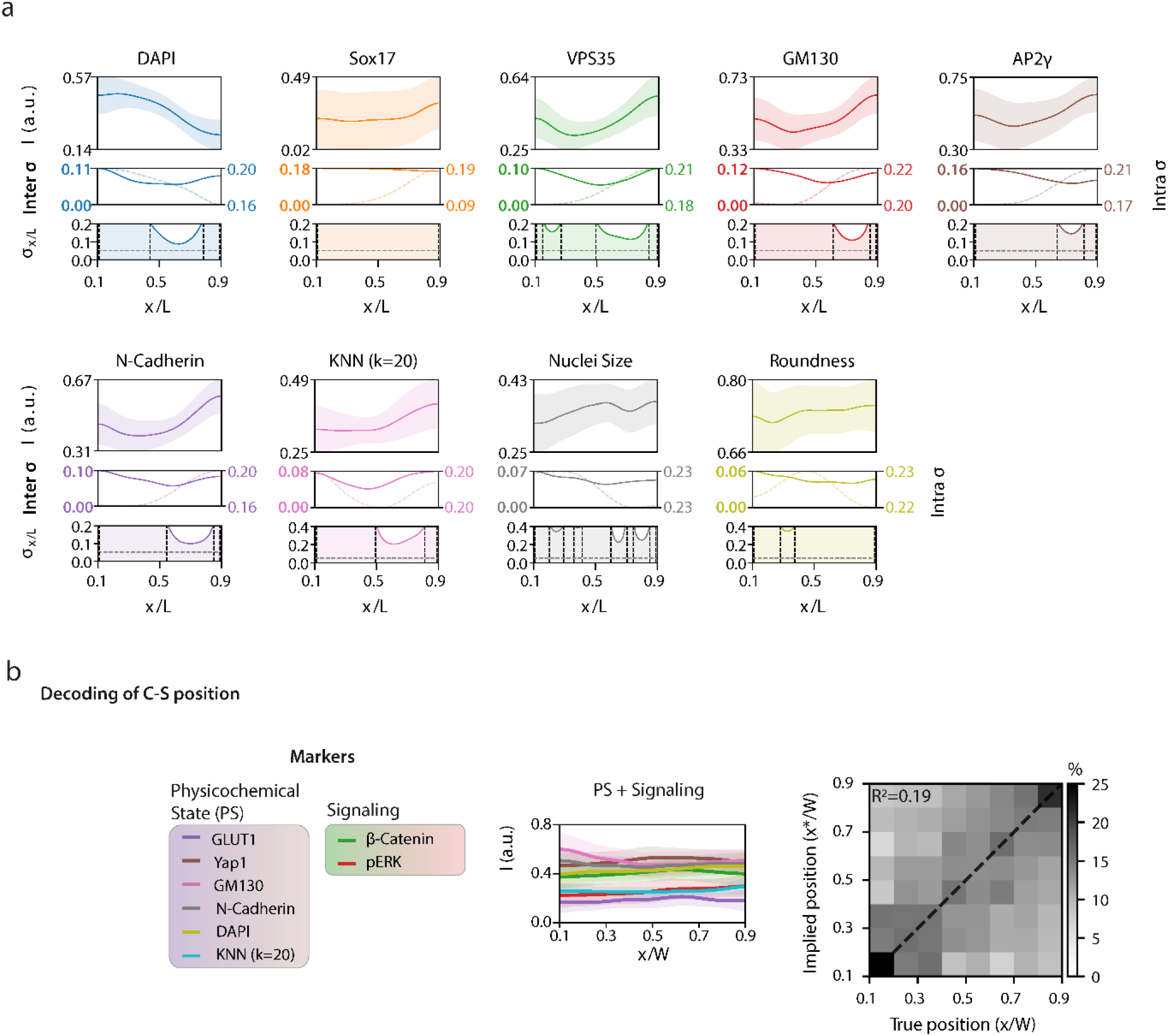
Spatial patterns of fate and cellular physicochemical states in 3D gastruloids and Bayesian decoding along the center-surface axis. a. Spatial profiles of multiplexed molecular, cellular, and morphological features along the normalized anterior-posterior (AP) axis of gastruloids (*x*/*L*; n=42). Profiles are shown for DAPI, Sox17, VPS35, GM130, AP2γ, N-Cadherin, KNN-based local density (*k* = 20), nuclear size, and nuclear roundness. Plots as in Fig.5c: Top rows show mean feature profiles, with bold lines indicating the average across gastruloids and shaded regions indicating the standard deviation. The middle row shows inter-gastruloid variability as continuous lines and in bold and intra-gastruloid variability as dotted lines. The bottom row shows positional error (σ_*x*/*L*_), quantifying the uncertainty in inferred cell position along the AP axis based on individual feature profiles. Horizontal dashed line corresponds to two effective nuclei diameters, vertical dashed lines to where positional error reaches 0.2, bounding zones with low positional error. b. Decoding of center-surface (CS) position with multimodal measurements in gastruloids. Middle: Mean intensity profiles (bold lines) along the normalized CS axis (*x*/*W*), with shaded regions indicating the standard deviation across gastruloids. Right: Decoding map *P*(*x*^*^|*x*) averaged across gastruloids. n=42. PS: Physicochemical State.

**supplementary Fig. 4.**
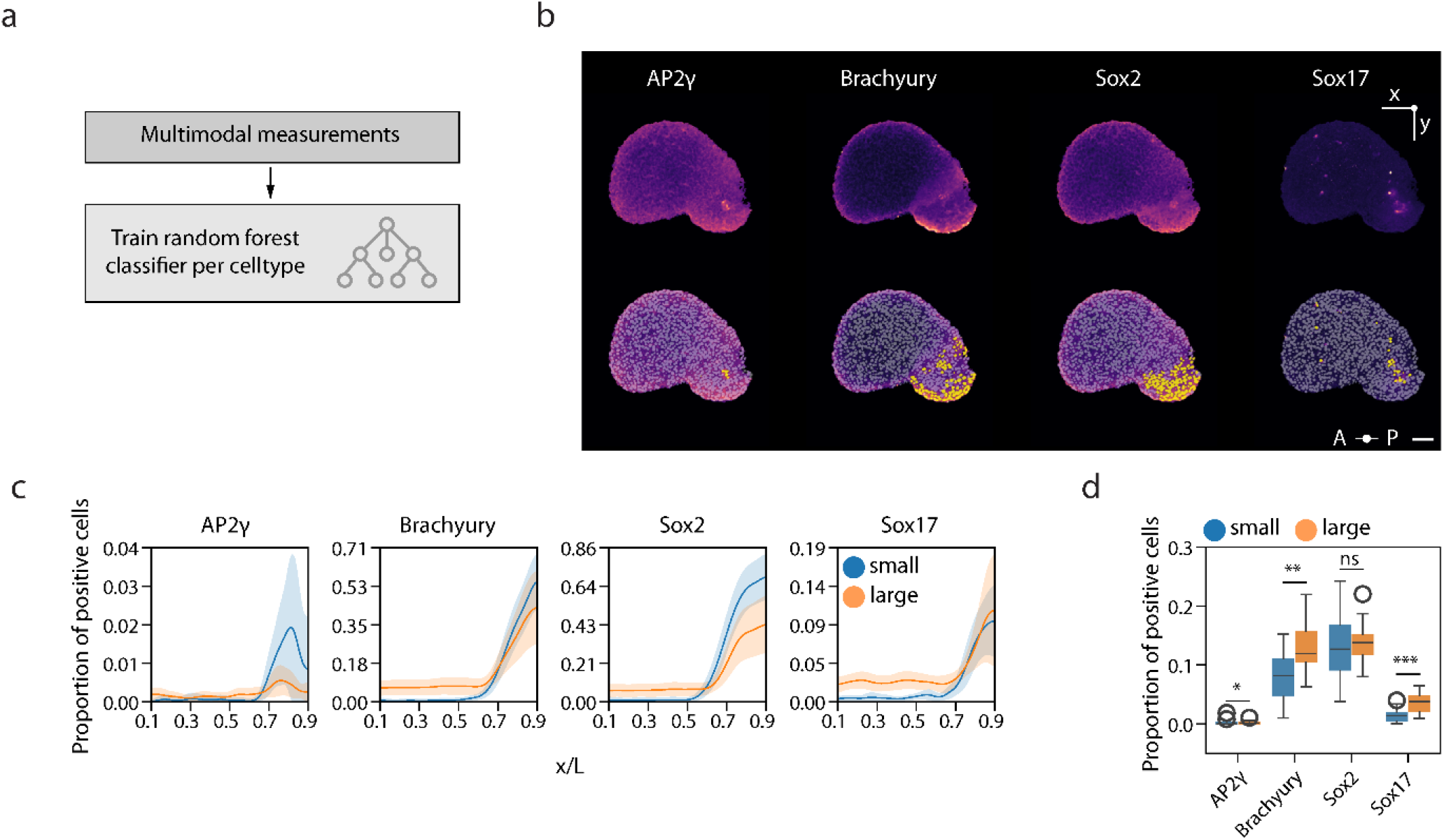
Cell type classification, spatial patterning, and lineage composition in gastruloids. a. Schematic of the random forest classifier workflow used for cell type classification. b. Representative 2D-slices of a gastruloid stained for AP2γ, Brachyury, Sox2, and Sox17, with corresponding cell type classification masks overlaid below. Yellow labels indicate cells classified as positive for the respective lineage marker. Scale bar 50 µm. c. Fraction of positive cells along the normalized AP axis (*x*/*L*) in small and large gastruloids (binned by nuclei count; blue and orange). Line indicates mean across gastruloids (n=42) and shading 95% Confidence Interval. d. Proportion of cells classified as positive for AP2γ, Brachyury, Sox2, or Sox17 in small (blue) and large (orange) gastruloids (n=42). Boxes, IQR with median; whiskers, 1.5x IQR; circles, outliers. Two-sided Mann-Whitney U test, small vs. large gastruloids (ns, not significant; * p < 0.05; ** p<0.01;*** p<0.001).

## References

1. Liberali, P. & Schier, A. F. The evolution of developmental biology through conceptual and technological revolutions. Cell 187, 3461–3495 (2024).

2. Hannezo, E. & Heisenberg, C.-P. Mechanochemical Feedback Loops in Development and Disease. Cell 178, 12–25 (2019).

3. Petridou, N. I., Grigolon, S., Salbreux, G., Hannezo, E. & Heisenberg, C.-P. Fluidization-mediated tissue spreading by mitotic cell rounding and non-canonical Wnt signalling. Nat Cell Biol 21, 169–178 (2019).

4. Dumortier, J. G. et al. Hydraulic fracturing and active coarsening position the lumen of the mouse blastocyst. Science 365, 465–468 (2019).

5. Martínez-Ara, G. et al. Optogenetic control of apical constriction induces synthetic morphogenesis in mammalian tissues. Nat Commun 13, 5400 (2022).

6. Palmquist, K. H. et al. Reciprocal cell-ECM dynamics generate supracellular fluidity underlying spontaneous follicle patterning. Cell 185, 1960–1973.e11 (2022).

7. Chan, C. J. et al. Hydraulic control of mammalian embryo size and cell fate. Nature 571, 112–116 (2019).

8. Shyer, A. E., Huycke, T. R., Lee, C., Mahadevan, L. & Tabin, C. J. Bending Gradients: How the Intestinal Stem Cell Gets Its Home. Cell 161, 569–580 (2015).

9. Kramer, B. A., Sarabia del Castillo, J. & Pelkmans, L. Multimodal perception links cellular state to decision-making in single cells. Science 377, 642–648 (2022).

10. Quail, D. F. & Walsh, L. A. Revolutionizing cancer research with spatial proteomics and visual intelligence. Nat Methods 21, 2216–2219 (2024).

11. Bodenmiller, B. Highly multiplexed imaging in the omics era: understanding tissue structures in health and disease. Nat Methods 21, 2209–2211 (2024).

12. Gut, G., Herrmann, M. D. & Pelkmans, L. Multiplexed protein maps link subcellular organization to cellular states. Science 361, eaar7042 (2018).

13. Black, S. et al. CODEX multiplexed tissue imaging with DNA-conjugated antibodies. Nat Protoc 16, 3802–3835 (2021).

14. Radtke, A. J. et al. IBEX: an iterative immunolabeling and chemical bleaching method for high-content imaging of diverse tissues. Nat Protoc 17, 378–401 (2022).

15. Hess, M. et al. Multiplexed embryo profiling links cellular state to zygotic genome activation in single cells. Preprint at 10.1101/2025.10.27.684723 (2025).

16. Tomer, R., Ye, L., Hsueh, B. & Deisseroth, K. Advanced CLARITY for rapid and high-resolution imaging of intact tissues. Nat Protoc 9, 1682–1697 (2014).

17. Choi, S. W., Guan, W. & Chung, K. Basic principles of hydrogel-based tissue transformation technologies and their applications. Cell 184, 4115–4136 (2021).

18. Yang, B. et al. Single-Cell Phenotyping within Transparent Intact Tissue through Whole-Body Clearing. Cell 158, 945–958 (2014).

19. Shah, S. et al. Single-molecule RNA detection at depth by hybridization chain reaction and tissue hydrogel embedding and clearing. Development 143, 2862–2867 (2016).

20. Park, Y.-G. et al. Protection of tissue physicochemical properties using polyfunctional crosslinkers. Nat Biotechnol 37, 73–83 (2019).

21. Lüthi, J. et al. Fractal: Towards FAIR bioimage analysis at scale with OME-Zarr-native workflows. Preprint at 10.64898/2026.03.05.709921 (2026).

22. Moore, J. et al. OME-Zarr: a cloud-optimized bioimaging file format with international community support. Histochem Cell Biol 160, 223–251 (2023).

23. Wilkinson, M. D. et al. The FAIR Guiding Principles for scientific data management and stewardship. Sci Data 3, 160018 (2016).

24. Marstal, K., Berendsen, F., Staring, M. & Klein, S. SimpleElastix: A User-Friendly, Multi-lingual Library for Medical Image Registration. in 2016 IEEE Conference on Computer Vision and Pattern Recognition Workshops (CVPRW) 574–582 (IEEE, Las Vegas, NV, USA, 2016). doi:10.1109/CVPRW.2016.78.

25. Hoffmann, M., Henninger, J., Veith, J., Richter, L. & Judkewitz, B. Blazed oblique plane microscopy reveals scale-invariant inference of brain-wide population activity. Nat Commun 14, 8019 (2023).

26. Pinheiro, D. & Heisenberg, C.-P. Zebrafish gastrulation: Putting fate in motion. in Current Topics in Developmental Biology vol. 136 343–375 (Elsevier, 2020).

27. Beccari, L. et al. Multi-axial self-organization properties of mouse embryonic stem cells into gastruloids. Nature 562, 272–276 (2018).

28. Van Den Brink, S. C. et al. Single-cell and spatial transcriptomics reveal somitogenesis in gastruloids. Nature 582, 405–409 (2020).

29. Wong, F. C. K. et al. NANOG is repurposed after implantation to repress Sox2 and begin pluripotency extinction. EMBO J 44, 5337–5374 (2025).

30. Ang, S.-L. et al. A targeted mouse *Otx2* mutation leads to severe defects in gastrulation and formation of axial mesoderm and to deletion of rostral brain. Development 122, 243–252 (1996).

31. Suppinger, S. et al. Multimodal characterization of murine gastruloid development. Cell Stem Cell 30, 867–884.e11 (2023).

32. Pour, M. et al. Emergence and patterning dynamics of mouse-definitive endoderm. iScience 25, 103556 (2022).

33. Cooke, C. B., Barrington, C., Baillie-Benson, P., Nichols, J. & Moris, N. Gastruloid-derived primordial germ cell-like cells develop dynamically within integrated tissues. Development 150, dev201790 (2023).

34. Hoshino, H., Shioi, G. & Aizawa, S. AVE protein expression and visceral endoderm cell behavior during anterior–posterior axis formation in mouse embryos: Asymmetry in OTX2 and DKK1 expression. Developmental Biology 402, 175–191 (2015).

35. Auman, H. J. et al. Transcription factor AP-2γ is essential in the extra-embryonic lineages for early postimplantation development. Development 129, 2733–2747 (2002).

36. Signaling regulation during gastrulation: Insights from mouse embryos and in vitro systems. in Current Topics in Developmental Biology vol. 137 391–431 (Elsevier, 2020).

37. Chhabra, S., Liu, L., Goh, R., Kong, X. & Warmflash, A. Dissecting the dynamics of signaling events in the BMP, WNT, and NODAL cascade during self-organized fate patterning in human gastruloids. PLoS Biol 17, e3000498 (2019).

38. Haantjes, R. R. et al. Towards an integrated view and understanding of embryonic signalling during murine gastrulation. Cells & Development 184, 204028 (2025).

39. Stapornwongkul, K. S. et al. Glycolytic activity instructs germ layer proportions through regulation of Nodal and Wnt signaling. Cell Stem Cell 32, 744–758.e7 (2025).

40. Schwayer, C. et al. Mechanosensation of Tight Junctions Depends on ZO-1 Phase Separation and Flow. Cell 179, 937–952.e18 (2019).

41. Scheibner, K. et al. Epithelial cell plasticity drives endoderm formation during gastrulation. Nat Cell Biol 23, 692–703 (2021).

42. Ozguldez, H. O. et al. Polarity inversion reorganizes the stem cell compartment of the trophoblast lineage. Cell Reports 42, 112313 (2023).

43. Freyer, L. et al. A loss-of-function and H2B-Venus transcriptional reporter allele for Gata6 in mice. BMC Dev Biol 15, 38 (2015).

44. Ichikawa, T. et al. Boundary-guided cell alignment drives mouse epiblast maturation. Nat. Phys. 22, 461–473 (2026).

45. Arnold, S. J. & Robertson, E. J. Making a commitment: cell lineage allocation and axis patterning in the early mouse embryo. Nat Rev Mol Cell Biol 10, 91–103 (2009).

46. Dingare, C., Cao, D., Yang, J. J., Sozen, B. & Steventon, B. Mannose controls mesoderm specification and symmetry breaking in mouse gastruloids. Developmental Cell 59, 1523–1537.e6 (2024).

47. Schindelin, J. et al. Fiji: an open-source platform for biological-image analysis. Nat Methods 9, 676–682 (2012).

48. Sofroniew, N. et al. napari: a multi-dimensional image viewer for Python. Zenodo 10.5281/ZENODO.14719463 (2025).

49. Manz, T. et al. Viv: multiscale visualization of high-resolution multiplexed bioimaging data on the web. Nat Methods 19, 515–516 (2022).

50. Peng, T. et al. A BaSiC tool for background and shading correction of optical microscopy images. Nat Commun 8, 14836 (2017).

51. Berg, S. et al. ilastik: interactive machine learning for (bio)image analysis. Nat Methods 16, 1226–1232 (2019).

52. Stringer, C., Wang, T., Michaelos, M. & Pachitariu, M. Cellpose: a generalist algorithm for cellular segmentation. Nat Methods 18, 100–106 (2021).

53. Sommer, C., Straehle, C., Kothe, U. & Hamprecht, F. A. Ilastik: Interactive learning and segmentation toolkit. in 2011 IEEE International Symposium on Biomedical Imaging: From Nano to Macro 230–233 (IEEE, Chicago, IL, USA, 2011). doi:10.1109/ISBI.2011.5872394.

54. Weigert, M. & Schmidt, U. Nuclei Instance Segmentation and Classification in Histopathology Images with Stardist. in 2022 IEEE International Symposium on Biomedical Imaging Challenges (ISBIC) 1–4 (IEEE, Kolkata, India, 2022). doi:10.1109/ISBIC56247.2022.9854534.

55. Wolny, A. et al. Accurate and versatile 3D segmentation of plant tissues at cellular resolution. eLife 9, e57613 (2020).

56. Hashmi, A. et al. Cell-state transitions and collective cell movement generate an endoderm-like region in gastruloids. eLife 11, e59371 (2022).

57. Mayran, A. et al. Cadherins modulate the self-organizing potential of pseudo-embryos. Cell Reports 44, 116567 (2025).

58. Underhill, E. J. & Toettcher, J. E. Control of gastruloid patterning and morphogenesis by the Erk and Akt signaling pathways. Development 150, dev201663 (2023).

59. Merle, M., Friedman, L., Chureau, C. & Gregor, T. Precise and scalable self-organization in mammalian pseudo-embryos. https://doi.org/10.48550/ARXIV.2303.17522 (2023) doi:10.48550/ARXIV.2303.17522.

60. Farag, N., Schiff, C. & Nachman, I. Coordination between Endoderm Progression and Gastruloid Elongation Controls Endodermal Morphotype Choice. http://biorxiv.org/lookup/doi/10.1101/2023.02.07.527329 (2023) doi:10.1101/2023.02.07.527329.

61. Technau, U. Brachyury, the blastopore and the evolution of the mesoderm. BioEssays 23, 788–794 (2001).

62. Anlaş, K. et al. Early autonomous patterning of the anteroposterior axis in gastruloids. Development 151, dev202171 (2024).

63. Kuhl, F. P. & Giardina, C. R. Elliptic Fourier features of a closed contour. Computer Graphics and Image Processing 18, 236–258 (1982).

64. Abraham, E. et al. YAP levels regulate anteroposterior elongation of hESC-derived gastruloids. Preprint at 10.64898/2025.12.01.691265 (2025).

65. Čapek, D. et al. EmbryoNet: using deep learning to link embryonic phenotypes to signaling pathways. Nat Methods https://doi.org/10.1038/s41592-023-01873-4 (2023) doi:10.1038/s41592-023-01873-4.

66. Bailleul, R. et al. Deciphering mechanical determinants of morphological evolution. Cell 189, 2598–2611.e18 (2026).

67. Lehr, S. et al. Self-Organised Pattern Formation in the Developing Neural Tube by a Temporal Relay of BMP Signalling. http://biorxiv.org/lookup/doi/10.1101/2023.11.15.567070 (2023) doi:10.1101/2023.11.15.567070.

68. Snijder, B. et al. Single-cell analysis of population context advances RNAi screening at multiple levels. Mol Syst Biol 8, MSB20129 (2012).

69. Mullins, M. C., Hammerschmidt, M., Haffter, P. & Nüsslein-Volhard, C. Large-scale mutagenesis in the zebrafish: in search of genes controlling development in a vertebrate. Current Biology 4, 189–202 (1994).

70. Wan, Y. et al. Whole-embryo spatial transcriptomics at subcellular resolution from gastrulation to organogenesis. Science 391, eadt3439 (2026).

71. Chang, T. et al. High-plex spatial RNA imaging in one round with conventional microscopes using color-intensity barcodes. Nat Biotechnol https://doi.org/10.1038/s41587-025-02883-7 (2025) doi:10.1038/s41587-025-02883-7.

72. Tkačik, G., Dubuis, J. O., Petkova, M. D. & Gregor, T. Positional Information, Positional Error, and Readout Precision in Morphogenesis: A Mathematical Framework. Genetics 199, 39–59 (2015).

73. Wolpert, L. Positional information and patterning revisited. Journal of Theoretical Biology 269, 359–365 (2011).

74. Tsai, T. Y.-C. et al. An adhesion code ensures robust pattern formation during tissue morphogenesis. Science 370, 113–116 (2020).

75. Lenne, P.-F. & Trivedi, V. Sculpting tissues by phase transitions. Nat Commun 13, 664 (2022).

76. Bragantini, J. et al. Ultrack: pushing the limits of cell tracking across biological scales. Nat Methods 22, 2423–2436 (2025).

77. Bunne, C. et al. Learning single-cell perturbation responses using neural optimal transport. Nat Methods 20, 1759–1768 (2023).

78. Klein, D. et al. CellFlow enables generative single-cell phenotype modeling with flow matching. Preprint at 10.1101/2025.04.11.648220 (2025).

79. Sun, X. et al. MIOFlow 2.0: A unified framework for inferring cellular stochastic dynamics from single cell and spatial transcriptomics data. Preprint at 10.48550/ARXIV.2603.22564 (2026).

80. Govindasamy, N. & Bedzhov, I. Isolation and Culture of Periimplantation and Early Postimplantation Mouse Embryos. in Comparative Embryo Culture (ed. Herrick, J. R.) vol. 2006 373–382 (Springer New York, New York, NY, 2019).

81. Rivera-Pérez, J. A., Diefes, H. & Magnuson, T. A simple enzymatic method for parietal yolk sac removal in early postimplantation mouse embryos. Developmental Dynamics 236, 489–493 (2007).

82. Smutny, M. et al. Friction forces position the neural anlage. Nat Cell Biol 19, 306–317 (2017).

83. Shamipour, S., Hofmann, L., Steccari, I., Kardos, R. & Heisenberg, C.-P. Yolk Granule Fusion and Microtubule Aster Formation Regulate Cortical Granule Translocation and Exocytosis in Zebrafish Oocytes. http://biorxiv.org/lookup/doi/10.1101/2022.08.10.503442 (2022) doi:10.1101/2022.08.10.503442.

84. Ichikawa, T. et al. An ex vivo system to study cellular dynamics underlying mouse peri-implantation development. Developmental Cell 57, 373–386.e9 (2022).

85. Chung, K. & Deisseroth, K. CLARITY for mapping the nervous system. Nat Methods 10, 508–513 (2013).

86. Gros, A. et al. A quantitative pipeline for whole-mount deep imaging and analysis of multi-layered organoids across scales. Preprint at 10.7554/eLife.107154.2 (2026).

87. McCormick, M., Liu, X., Jomier, J., Marion, C. & Ibanez, L. ITK: enabling reproducible research and open science. Front. Neuroinform. 8, (2014).

88. Savill, R. G. et al. SpinePy enables automated 3D spatiotemporal quantification of multicellular in vitro systems. Preprint at 10.1101/2025.09.10.674634 (2025).

89. Albergante, L. et al. Robust and Scalable Learning of Complex Intrinsic Dataset Geometry via ElPiGraph. Entropy 22, 296 (2020).

90. Mederacke, M. et al. The emergence of the fractal bronchial tree. Preprint at 10.1101/2025.01.13.632436 (2025).

91. Van Der Walt, S. et al. scikit-image: image processing in Python. PeerJ 2, e453 (2014).

92. Petkova, M. D., Tkačik, G., Bialek, W., Wieschaus, E. F. & Gregor, T. Optimal Decoding of Cellular Identities in a Genetic Network. Cell 176, 844–855.e15 (2019).

93. Kuhl, F. P. & Giardina, C. R. Elliptic Fourier features of a closed contour. Computer Graphics and Image Processing 18, 236–258 (1982).

